# A measles-vectored COVID-19 vaccine induces long-term immunity and protection from SARS-CoV-2 challenge in mice

**DOI:** 10.1101/2021.02.17.431630

**Authors:** Phanramphoei N. Frantz, Aleksandr Barinov, Claude Ruffié, Chantal Combredet, Valérie Najburg, Samaporn Teeravechyan, Anan Jongkaewwattana, Matthieu Prot, Laurine Conquet, Xavier Montagutelli, Priyanka Fernandes, Hélène Strick-Marchand, James Di Santo, Etienne Simon-Lorière, Christiane Gerke, Frédéric Tangy

## Abstract

In light of the expanding SARS-CoV-2 pandemic, developing efficient vaccines that can provide sufficient coverage for the world population is a global health priority. The measles virus (MV)-vectored vaccine is an attractive candidate given the measles vaccine’s extensive safety history, well-established manufacturing process, and induction of strong, long-lasting immunity. We developed an MV-based SARS-CoV-2 vaccine using either the full-length spike (S) or S2 subunit as the antigen. While the S2 antigen failed to induce neutralizing antibodies, the prefusion-stabilized, full-length S (MV-ATU2-SF-2P-dER) construct proved to be an attractive vaccine candidate, eliciting strong Th1-dominant T-cell and neutralizing antibody responses against the S antigen while minimizing reactivity to the vector itself. Neutralizing antibody titers remained high three months after homologous prime-boost immunization, and infectious virus was undetectable in all animals after challenge with a mouse-adapted SARS-CoV-2 virus.

## Introduction

More than 50 million people worldwide have been affected by the SARS-CoV-2 pandemic to date and more than 1.2 million have died of COVID-19, the disease caused by this newly emerged coronavirus. The pandemic has resulted in unprecedented global social and economic disruption, with a projected “optimistic loss” of $3.3 trillion and a worst-case scenario loss of $82 trillion worldwide ^1^. While the virus is now explosively expanding in a second wave in Europe and the northern hemisphere, no specific treatment has been shown to prevent or cure the disease. Together with enforcing public health measures, effective vaccines that can prevent infection, disease, and transmission, and their deployment on a global scale are required for a return to pre-COVID-19 normalcy.

An ideal vaccine against SARS-CoV-2 should be safe, able to induce long-lasting protective immune responses after a single administration, be produced at low cost and easily scaled up for mass vaccination. Numerous vaccine platforms are currently being used to develop SARS-CoV-2 vaccines ^2^. Among them, live attenuated viral vectors look particularly interesting as they induce lasting protective immunity after a single dose and are inexpensive to manufacture at large scale. In particular, the live attenuated measles vaccine (MV) is one of the safest and most efficacious human preventive medicines. It elicits neutralizing antibodies and robust, long-lasting Th1 cellular responses, making it an attractive candidate for SARS-CoV-2 vaccination with minimal risk of vaccine-associated enhanced respiratory disease (VAERD) ^3^.

SARS-CoV-2 is an enveloped single-stranded positive-sense RNA virus belonging to the *Coronavidae* family and the β-coronavirus genus ^4^. Whole genome sequencing of SARS-CoV-2 revealed 79.6% nucleotide sequence similarity with SARS-CoV-1 ^5^. The genome of SARS-CoV-2 encodes 4 structural proteins: the spike protein (S), the envelope protein (E), the membrane protein (M), and the nucleocapsid (N). The S protein, a trimeric class I fusion protein localized on the surface of the virion, plays a central role in viral attachment and entry into host cells. Cleavage of the S protein into S1 and S2 subunits by host proteases ^6^ is essential for viral infection. The S1 subunit contains the receptor-binding-domain (RBD), which enables the virus to bind to its entry receptor, the angiotensin-converting enzyme 2 (ACE2) ^4, 7^. After docking with the receptor, the S1 subunit is released and the S2 subunit reveals its fusion peptide to mediate membrane fusion and viral entry ^8^.

The coronavirus S protein contains the major epitopes targeted by neutralizing antibodies and is thus considered as a main antigen for developing vaccines against human coronaviruses ^8, 9, 10, 11, 12^. Antibodies targeting the RBD may neutralize virus by blocking viral binding to receptors on host cells and preventing entry. Additionally, it has been observed that synthetic peptides mimicking and antibodies targeting the second heptad region (HR2) in the S2 subunit of SARS-CoV have strong neutralizing activity ^13, 14, 15, 16, 17^, likely by preventing the conformational changes required for membrane fusion. Efforts to develop a SARS-CoV-2 vaccine have thus focused on eliciting responses against the S protein.

A number of recombinant MV (rMV)-based vaccines against viral pathogens are currently in preclinical and clinical trials ^18^. An rMV-based vaccine against chikungunya virus was demonstrated to be well-tolerated and immunogenic in phase I and II clinical trials, eliciting 90% seroconversion after a single immunization and 100% after boost despite the presence of preexisting measles immunity in volunteers ^19, 20^. Other MV-based candidates currently in clinical development include vaccines against Zika and Lassa viruses ^21, 22^. We also previously showed that rMV expressing the unmodified SARS-CoV-1 S protein induced a Th1-oriented response with high titers of neutralizing antibodies that protected immunized mice from infectious intranasal challenge by SARS-CoV-1 ^9^. The MV-MERS-CoV vaccine has also yielded promising preclinical results ^23^. Given the excellent safety and efficacy profiles of these vaccine candidates, an MV-based vaccine targeting the S protein of SARS-CoV-2 has great potential to be both safe and effective.

To explore this potential, we generated a series of rMVs expressing either full-length S or the S2 subunit protein of SARS-CoV-2 in prefusion-stabilized or native forms and tested their capacity to elicit neutralizing antibodies and T-cell responses. Additionally, we examined their ability to protect from SARS-CoV-2 challenge in a mouse model of measles vaccination.

## Results

### Design of SARS-CoV-2 S antigens

Based on our previous work with MV expressing SARS-CoV-1 S ^9^ and since SARS-CoV and SARS-CoV-2 S proteins share a high degree of similarity ^24^, the full-length S protein of SARS-CoV-2 was chosen as the main antigen to be expressed by the MV vector. We introduced a number of modifications in the native S sequence to improve its expression and immunogenicity (Fig. 1). First, the RNA sequence was codon-optimized to increase its expression in human cells. Second, we mutated two prolines, K986P and V987P, in the S2 region to generate a subset of 2P constructs, following a proven strategy to stabilize the S protein in its prefusion conformation, increasing its expression and immunogenicity ^25, 26, 27^. Third, to increase the surface expression of the S protein in MV-infected cells, we deleted the 11 C-terminal amino acids (aa 1263-1273) from the S cytoplasmic tail to generate dER constructs. The cytoplasmic tail of coronaviruses S proteins contains one or two distinct retention signals: the endoplasmic reticulum retrieval signal (ERRS) comprising KxHxx or KKxx motifs, and the tyrosine-dependent localization signal Yxxϕ ^28^. S proteins with ERRS are recruited into coatomer complex I (COPI) and recycled from the Golgi to the ER in retrograde. Thus, the repeated cycling of S proteins between the ER and the Golgi leads to S protein intracellular retention, while mutant S proteins lacking the ERRS are transported to the plasma membrane ^29,30^. Similarly, the S proteins of *Alphacoronaviruses* with the Yxxϕ motif are retained in the ER with little or no S protein trafficking to the cell surface ^31^. We therefore designed our SARS-CoV-2 S antigens with the deletion of all possible retention signals from the cytoplasmic tail.

**Fig 1.**
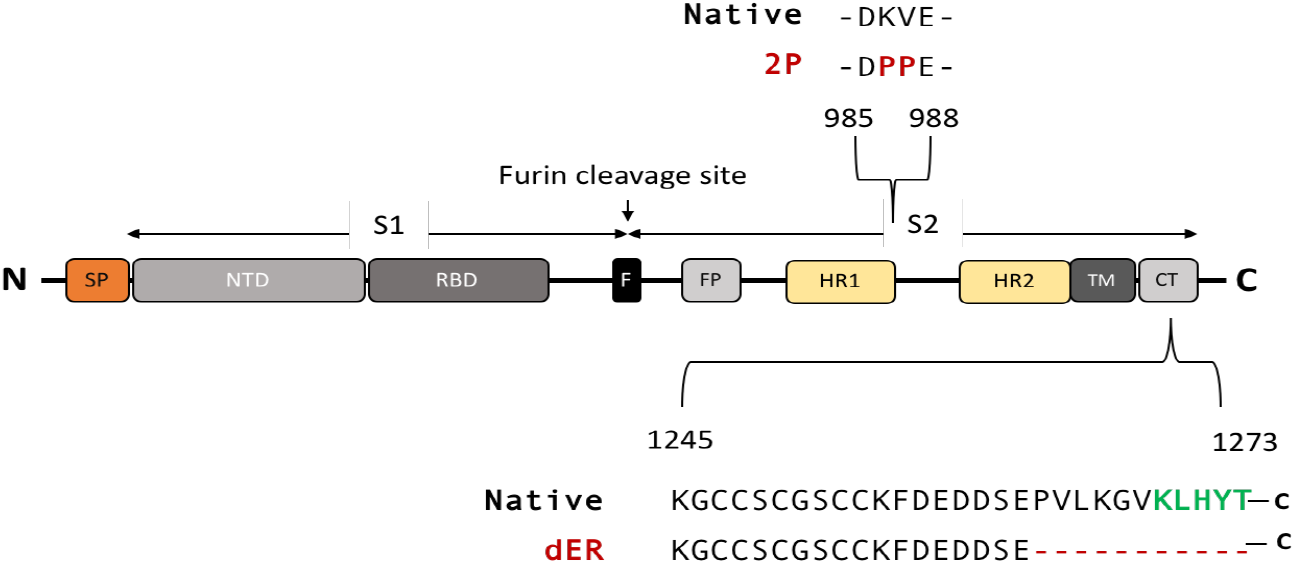
Schematic of the native S protein of SARS-CoV-2. The native S protein is 1273 amino acids (aa) in length. The protein contains 2 subunits, S1 and S2, generated by cleavage at the furin cleavage site (F). S1 contains the signal peptide (SP), N-terminal domain (NTD) and receptor-binding domain (RBD). S2 contains the fusion peptide (FP), heptad repeats 1 (HR1) and 2 (HR2), transmembrane domain (TM), and cytoplasmic tail (CT). The 2P indicates the two mutated prolines, K986P and V987P. The green letters indicate the endoplasmic reticulum retrieval signal (ERRS) motif, KxHxx, in the CT. dER indicates constructs carrying a deletion of the 11 C-terminal amino acids from the CT.

To investigate the possibility of generating a broad-spectrum vaccine targeting both SARS-CoV-1 and SARS-CoV-2 clinical isolates, we also designed S2 subunit antigens (Fig. 2a). The S2 subunit of SARS-CoV-2 is highly conserved among SARS-like CoVs and shares 99% identity with those of bat SARS-like CoVs (SL-CoV ZXC21 and ZC45) and of a human SARS-CoV-1 ^24^. We therefore designed S2 subunit antigens, both in its native trimer and prefusion-stabilized form, with the signal peptide of the S protein inserted in the N-terminus to target the antigen to the cell surface.

**Fig 2.**
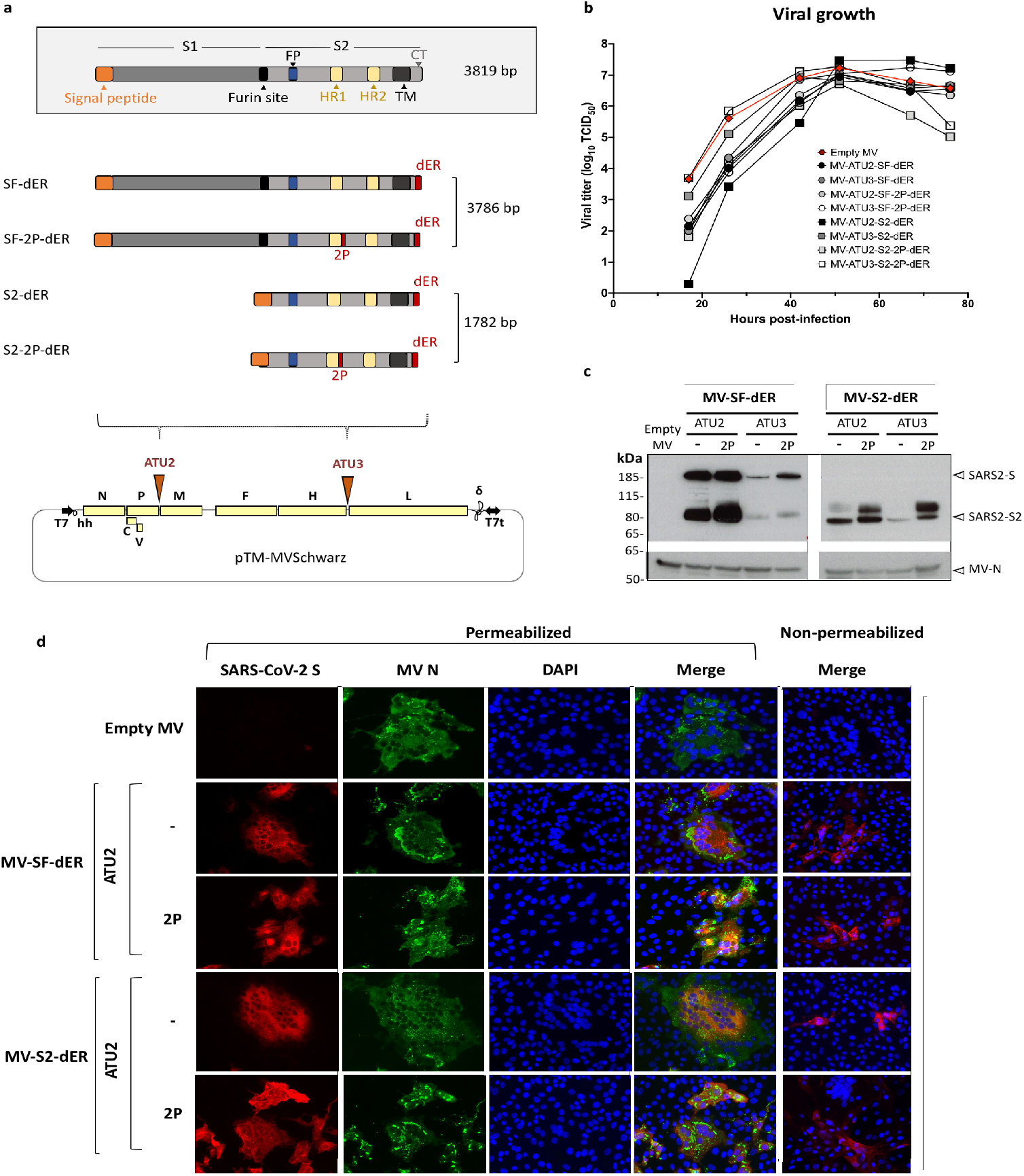
Schematic of S gene constructs and characterization of S-expressing rMVs. **a)** The native S gene of SARS-CoV-2 with notable domains indicated in color boxes relative to the S gene constructs cloned into the MV vector. 2P and dER modifications indicated by the red boxes. All S constructs were cloned into either the second (ATU2) or third (ATU3) additional transcription units of pTM-MVSchwarz (MV Schwarz), the MV vector plasmid. The MV genome comprises the nucleoprotein (N), phosphoprotein (P), V and C accessory proteins, matrix (M), fusion (F), hemagglutinin (H) and polymerase (L) genes. Plasmid elements include the T7 RNA polymerase promoter (T7), hammerhead ribozyme (hh), hepatitis delta virus ribozyme (∂), and T7 RNA polymerase terminator (T7t). **b)** Growth kinetics of rMV constructs used to infect Vero cells at an MOI of 0.1. Cell-associated virus titers are indicated in TCID_50_/ml. **c)** Western blot analysis of SARS-CoV-2 S protein in cell lysates of Vero cells infected with the rMVs expressing SF-dER or S2-dER from either ATU2 or ATU3, with or without the 2P mutation. **d)** Immunofluorescence staining of Vero cells infected with the indicated rMVs 24 h after infection. Permeabilized or non-permeabilized cells were stained for S (red), MV N (green) and nuclei (blue).

Altogether, we designed four different SARS-CoV-2 S constructs (Fig. 2a): 1) the native-conformation full-length S trimer (SF-dER); 2) the prefusion-stabilized full-length S (SF-2P-dER); 3) the native conformation trimer S2 subunit (S2-dER); and 4) the prefusion-stabilized S2 subunit (S2-2P-dER).

### Expression profile of SARS-CoV-2 S antigens

Full-length S and S2 sequences were firstly cloned into pCDNA and transfected into HEK293T cells to verify expression and assess surface protein localization by surface staining followed by flow cytometry. Prefusion-stabilized S constructs were observed to localize more strongly to the surface of transfected cells (S. Fig.1). Functionality of the S proteins was analyzed by transfecting the same pCDNA vectors in Vero cells, which express ACE-2. Once S proteins bound to ACE-2 receptors, activation of the fusion protein can be observed through the formation of large syncytia among cells. Vero cells expressing the native S protein (full-length S with an intact CT) exhibited significant syncytium formation (S. Fig.2), indicating that functional S proteins were expressed on the cell surface. Notably, the SdER mutants induced increased fusion compared to native SF, confirming the expected increased surface expression of the S protein when the ERRS is deleted. Interestingly, expression of the S2 subunit alone resulted in a hyper-fusion phenotype in Vero cells. This suggests the triggering of non-receptor-mediated membrane fusion by proteases cleaving at the S2’ site and freeing the fusion peptide. On the contrary, both the 2P-stabilized SF-2P-dER and S2-2P-dER did not induce syncytium formation, indicating that their fusion activity was abrogated by the 2P mutation.

### Generation of rMVs expressing SARS-CoV-2 S and S2 proteins

The four antigenic constructs were individually cloned into the pTM-MVSchwarz plasmid at additional transcription units (ATU), with ATU2 located between the P and M genes of the MV genome and ATU3 between the H and L genes ^32^ (Fig. 2a). Due to the decreasing expression gradient of MV genes cloning in ATU2 allows high-level expression of the antigen while cloning in ATU3 results in lower levels of expression ^33^. The lower expression from ATU3 is a trade-off to facilitate rescue of rMV encoding antigens that are toxic or difficult to express.

All rMVs expressing the S proteins were successfully rescued by reverse genetics and propagated in Vero cells. Although the rMVs exhibited slightly delayed growth kinetics, final virus yields were high and identical to that of the parental MV Schwarz (∼10 ^7^ TCID_50_/ml) (Fig. 2b). The expression of S antigens was detected in infected Vero cells by western blotting (WB) and immunofluorescence staining (IF) (Fig. 2c, d and S. Fig.3, 4). As expected, much higher antigen expression was observed from ATU2 vectors compared to ATU3 (Fig. 2c).

When using recombinant viral vectors as vaccines, the genetic stability of constructs is a major concern as it guarantees the effectiveness of the vaccine after multiple manufacturing steps. Such analysis of rMVs after serial passaging in Vero cells revealed that MV-ATU2-SF-dER, which expresses the native S from ATU2, was unstable, with loss of S expression by passage 5 (S. Fig.5). In contrast, its 2P counterpart was stably and efficiently expressed up to passage 10. The sequence of MV-ATU2-SF-dER genome was also confirmed by Next-Generation Sequencing (NGS) (S. Fig.6), with no mutations detected in the antigen. Therefore, we discarded the vaccine candidates expressing native S and selected those expressing the prefusion-stabilized SF-2P and S2-2P constructs for further immunogenicity studies.

### Induction of SARS-CoV-2 neutralizing antibodies in mice

We investigated the immunogenicity of selected rMV vaccine candidates in IFNAR ^−/−^ mice susceptible to MV infection ^34^. Animals were immunized by one or two intraperitoneal administrations of the rMV candidates at 1×10 ^5^ TCID_50_ on days 0 and 30 (Fig. 3a). Empty MV Schwarz was used for control vaccination. Mice sera were collected 4 weeks after the prime and 12 days after the boost. The presence of S-and MV-specific IgG antibodies was assessed by indirect ELISA using SARS-CoV-2 S recombinant protein and native MV antigens, respectively.

**Fig 3.**
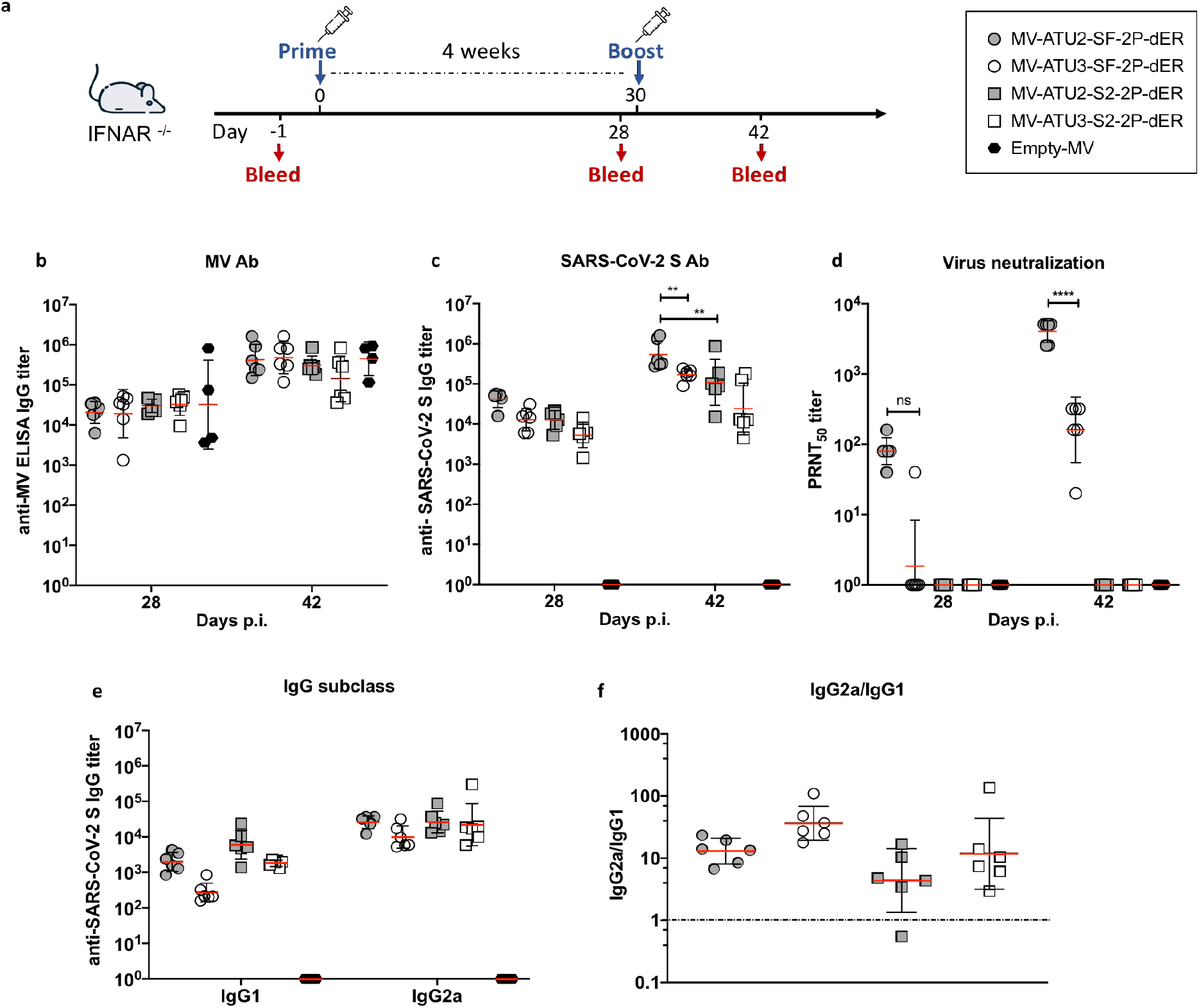
Induction of humoral responses by prime-boost vaccination. **a)** Homologous prime-boost of IFNAR −/−mice (*n*=6 or *n*=4 for the empty MV control) immunized intraperitoneally with 1×10 ^5^ TCID_50_ of the indicated rMV at days 0 and 28. Sera were collected 28 and 42 days after immunization and assessed for specific antibody responses to **b)** MV antigens or **c)** S-SARS-CoV-2 S. The data show the reciprocal endpoint dilution titers with each data point representing an individual animal. **d)** Neutralizing antibody responses to SARS-CoV-2 virus expressed as 50% plaque reduction neutralization test (PRNT_50_) titers. **e)** IgG subclass of S-specific antibody responses in mice 4 weeks after the first immunization. **f)** Ratio of IgG2a/IgG1 or Th1/Th2 responses. Data are represented as geometric mean with line and error bars indicating geometric SD. Statistical significance was determined by a two-way ANOVA adjusted for multiple comparisons. Asterisks (*) indicate significant mean differences (** *p*<0.01, and **** *p*<0.001) as determined by the Mann-Whitney *U*-test.

All animals raised high MV-specific IgG antibodies after prime at comparable titers in all groups (∼10 ^4^–10 ^5^ IgG titer), indicating efficient vaccine take in all the animals (Fig. 3b). Boost immunization increased MV-specific antibody titers in all groups, indicating that all animals received a successful prime-boost vaccination. Specific IgG antibodies to SARS-CoV-2 S were detected in 100% of immunized mice. Interestingly, rMVs expressing SF-2P-dER or S2-2P-dER antigens from ATU2 elicited higher levels of anti-S antibodies than the ATU3 vectors, particularly after boosting (Fig. 3c). Pre-immune sera and sera from control animals that received empty MV remained negative for anti-S antibodies (data not shown).

We next assessed the presence of SARS-CoV-2 neutralizing antibodies (NAbs) using plaque reduction neutralization tests (PRNT) with SARS-CoV-2 virus infection of Vero E6 monolayers. After the prime, SARS-CoV-2 NAbs were found in all mice immunized with SF-2P-dER expressed from ATU2 but only one mouse immunized with the ATU3 construct (Fig. 3d). After the second immunization, NAb titers increased in both groups, with the ATU2 group exhibiting ten-fold higher NAb titers compared to the ATU3 group. No NAbs were detected in animals immunized with the S2 candidates despite the high levels of anti-S antibodies (Fig. 3c).

As IgG isotype switching can serve as indirect indicators of Th1 and Th2 responses ^35^, we determined S-specific IgG1 and IgG2a isotype titers in the sera of immunized mice (Fig. 3f, g). Similar to our previous results ^9^, rMV candidates elicited significantly higher IgG2a antibody titers than IgG1, reflecting a predominant Th1-type immune response (Fig. 3f, g). Since activated T cells play important roles in shaping Th1 and Th2 cytokine production, we analyzed S-specific T-cell responses in MV-immunized mice in more detail.

### Induction of S-specific T-cell responses

Cell-mediated immune responses elicited by immunization were first investigated using an IFN-γ ELISPOT assay. Groups of IFNAR ^−/−^ mice were sacrificed one week after prime immunization (Fig. 4a). To evaluate S-specific responses, splenocytes were stimulated *ex vivo* with a pool of synthetic peptides covering the predicted CD8 ^+^ and CD4 ^+^ T-cell epitopes of the SARS-CoV-2 S protein, matching the MHC-I H-2K ^b^/H-2D ^b^ and MHC-II I-A ^b^ haplotype of 129sv IFNAR ^−/−^ mice (S.Table 1). Splenocytes were also stimulated with an empty MV virus to detect MV vector-specific T-cell responses.

**Fig 4.**
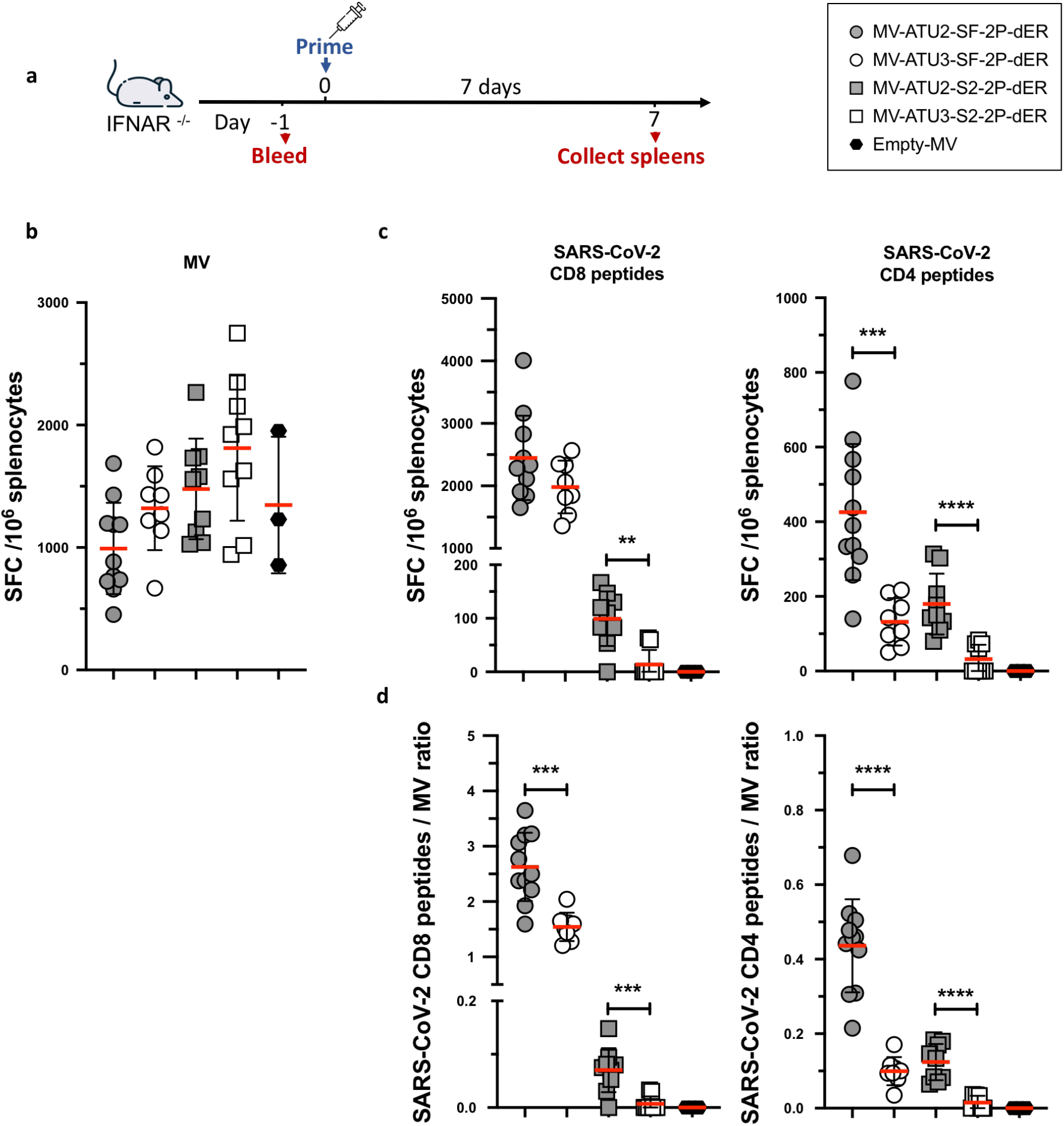
Induction of S-specific cellular responses by rMV vaccination. **a)** Immunization of IFNAR −/−mice (*n*=12 or *n*=3 for the empty MV control) immunized intraperitoneally with 1×10 ^5^ TCID_50_ of the indicated rMVs. Seven days after immunization, ELISPOT for IFNγ was performed on freshly extracted splenocytes. The data are shown as IFNγ-secreting cells or spot-forming cells (SFC) per 1×10 ^6^ splenocytes detected after stimulating with **b)** MV Schwarz or **c)** SARS-CoV-2 S peptide pools specific to CD8 ^+^ or CD4 ^+^ T cells. **d)** Ratio of IFNγ-secreting cells stimulated by CD4 ^+^ or CD8 ^+^ peptides to those stimulated by MV Schwarz. Each data point represents an individual mouse. Asterisks (*) indicate significant mean differences (* *p*<0.05; ** *p*<0.01, and **** *p*<0.001) as determined by the Mann-Whitney *U*-test.

High levels of T-cell responses to SARS-CoV-2 S and MV were elicited early after prime vaccination (Fig. 4). Splenocytes from mice vaccinated with MV-ATU2-SF-2P-dER yielded remarkably high IFN-γ secretion levels after stimulation with an MHC class I-restricted S peptide pool, yielding around 2,500 spot forming cells (SFC) per 10 ^6^ splenocytes. Lower IFN-γ responses were observed upon stimulation with MHC class II-restricted S peptides, at approximately 400 SFC/10 ^6^ splenocytes (Fig. 4b, c). Splenocytes of these mice also exhibited relatively low vector-specific IFN-γ responses (∼990 SFC/10 ^6^ splenocytes), indicating a well-balanced S-to-MV vector response ratio (Fig. 4d). The ATU3 counterpart of the same vaccine tended to generate more vector-specific IFN-γ secreting cells (∼1,320 SFC/10 ^6^ splenocytes), while at the same time being less efficient in producing S-specific IFN-γ secreting cells after stimulation with MHC class II-restricted S peptides (∼130 SFC/10 ^6^ splenocytes). While IFN-γ responses after stimulation with MHC class I-restricted S peptides were not significantly different from those of its ATU2 counterpart (∼1,980 SFC/10 ^6^ splenocytes), the S-to-MV vector response ratio was significantly higher for MV-ATU2-SF-2P-dER (Fig. 4b).

In contrast, S-specific IFN-γ responses elicited by S2-only constructs were low after stimulation with either of the S peptide pools (Fig. 4c, d). S-to-MV vector response ratios also remained very low in these animals, suggesting that immunization with the S2 protein subunit alone might be not sufficient to induce strong protective cellular immune responses (Fig. 4e, f). MV-ATU3-S2-2P-dER and empty MV were unable to induce S-specific IFN-γ responses.

We next studied S-specific CD4 ^+^ and CD8 ^+^ T cells by flow cytometric analysis after intracellular cytokine staining (ICS). S-specific IFN-γ ^+^ and TNF-α ^+^ responses were observed in for CD8 ^+^ T cells, while CD4 ^+^ T cells responded poorly to S peptide pool stimulation (Fig. 5a, b). Similar to the ELISPOT results, SF-2P-dER expressed from ATU2 or ATU3 induced remarkably high and comparable percentages of S-specific IFN-γ ^+^ and TNF-α ^+^ CD8 ^+^ T cells, while the S2 protein expressed from ATU2 was ten times less immunogenic. MV-ATU3-S2-2P-dER and empty MV were unable to induce S-specific IFN-γ-or TNF-α-producing T-cells. IL-5-secreting cells (indicative of a Th2-biased response) were not detected in any of the immunization groups. An additional detailed analysis of T cell responses in mice immunized with MV-ATU2-SF-2P-dER confirmed the strong stimulation of CD8 compartment with high levels of S-specific IFN-γ ^+^ and TNF-α ^+^ producing CD8 ^+^ T cells, as well as double positive IFN-γ ^+^ / TNF-α ^+^ producing CD8 ^+^ T cells. No IL-5 or IL-13 was detected in CD4 ^+^ or CD8 ^+^ T cells, as well as in CD4 ^+^ / CD44 ^+^ / CD62L ^-^ memory T cells, confirming that S-specific memory T cells are also Th1-oriented (S. Fig. 7).

**Fig 5.**
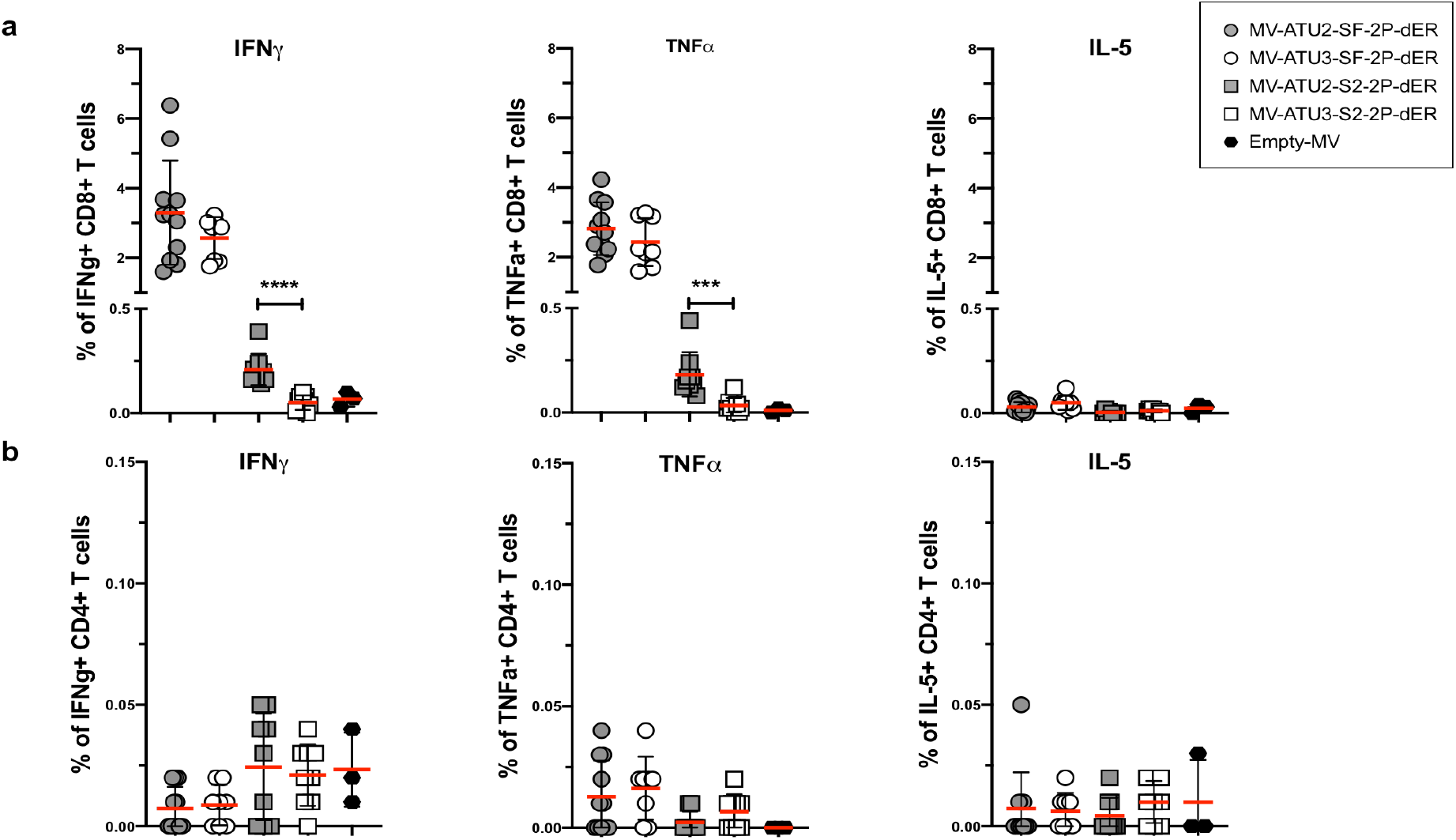
Cytokine expression profile of T cells. IFNAR −/−mice (*n*=12 or *n*=3 for the empty MV control) immunized intraperitoneally (i.p.) with 1×10 ^5^ TCID_50_ of the indicated rMVs and splenocytes were stimulated with S-specific peptide pools. S-specific **a)** CD8 ^+^ and **b)** CD4 ^+^ T-cells were stained for intracellular IFNγ, TNFα and IL-5. Asterisks (*) indicate significant mean differences (* *p*<0.05; ** *p*<0.01, and **** *p*<0.001) as determined by the Mann-Whitney *U*-test.

Taken together, these results demonstrate that MV-ATU2-SF-2P-dER induces a robust Th1-driven T-cell immune response to SARS-CoV-2 S antigens at significantly higher levels than MV-ATU3-SF-2P-dER. The S2 candidates elicited much lower cellular responses, as observed previously with NAb levels, indicating that S2 alone is not sufficient to induce an efficient immune response in these mice. We therefore excluded the S2 candidates from further analysis.

### Persistence of neutralizing antibodies and protection from intranasal challenge

We monitored the persistence of anti-S antibodies in mice immunized twice with either MV-ATU2-SF-2P-dER or MV-ATU3-2F-2P-dER (Fig. 6a). As usually observed for MV responses (Fig. 6b), S-specific IgG titers persisted and stabilized at high levels (10 ^5^–10 ^6^ limiting dilution titers) for both ATU2 and ATU3 candidates for up to three months after boosting (Fig 6c). However, immunization with the ATU2 construct resulted in significantly higher levels of S-specific IgG and NAb titers (10 ^3^–10 ^4^ limiting dilution titers) over the duration of the experiment (Fig. 6d).

**Fig 6.**
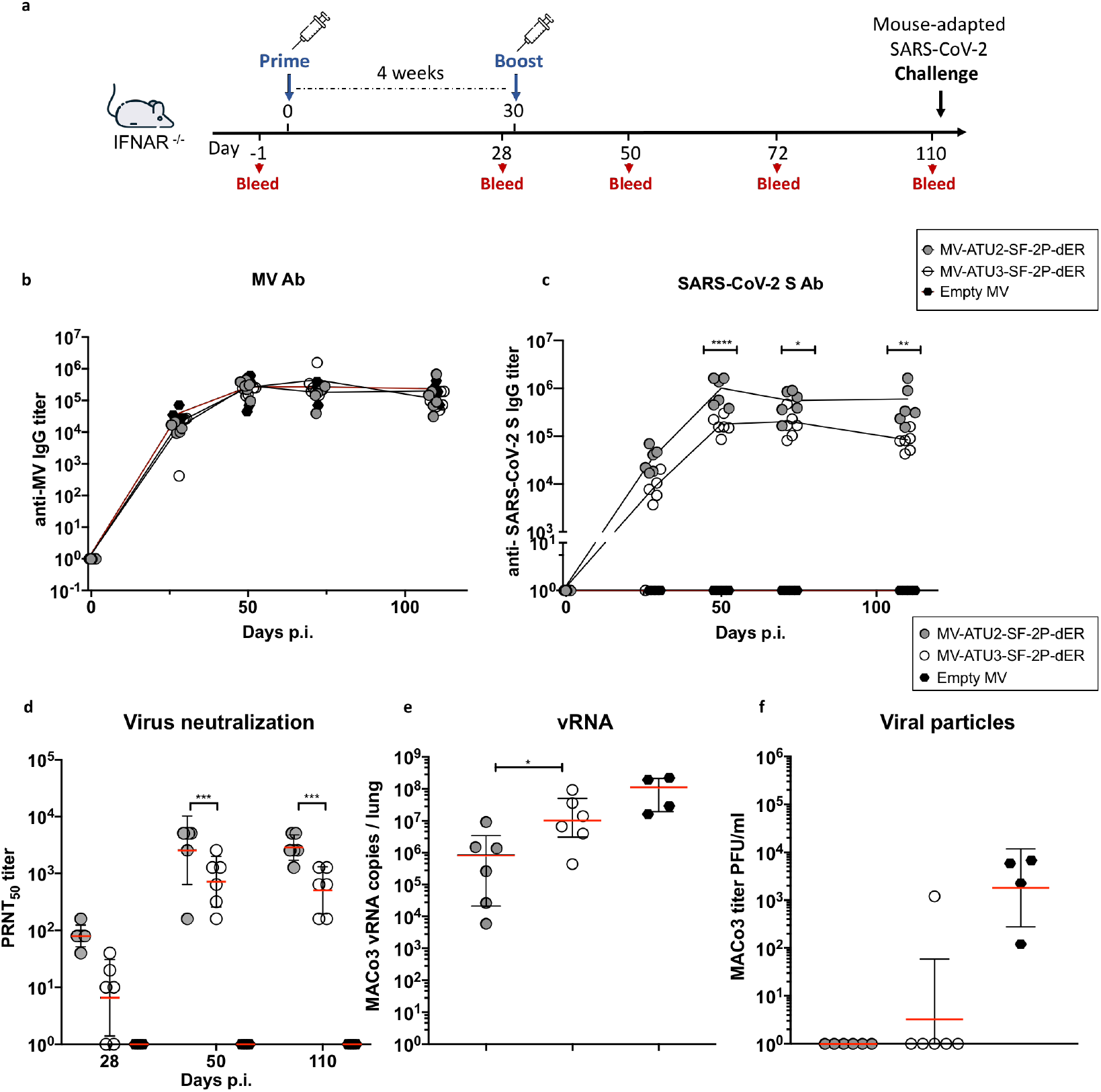
Persistence of neutralizing antibodies and immune protection. **a)** Immunization and challenge schedule for IFNAR −/−mice (*n*=6). Animals were immunized interperitoneally by homologous prime-boost at days 0 and 28. Sera were collected at days 52, 72, and 110. Animals were challenged on day 110 by intranasal inoculation of mouse-adapted SARS-CoV-2 virus (MACo3) at 1.5 x 10 ^5^ PFU. Sera were assessed for levels of specific antibodies against **b)** MV and **c)** SARS-CoV-2 S. **d)** Neutralizing antibody responses against SARS-CoV-2 virus, expressed as 50% plaque reduction neutralization test (PRNT_50_) titers. **e)** SARS-CoV-2 viral RNA copies detected by RT-qPCR in homogenized lungs of challenged animals, calculated as copies/lung. **f)** Titer of infectious viral particles recovered from the homogenized lung of the immunized animals expressed as PFU/lung. Data are represented as geometric means with line and error bars indicating geometric SD. Statistical significance was determined by a two-way ANOVA adjusted for multiple comparisons. Asterisks (*) indicate significant mean differences (* *p*<0.05; ** *p*<0.01, and **** *p*<0.001).

To determine whether these responses confer protection from SARS-CoV-2 infection, immunized mice were challenged intranasally with 1.5 x 10 ^5^ PFU of MACo3, a mouse-adapted SARS-CoV-2 virus (Montagutelli et al, in prep). Three days after challenge, mice were sacrificed and the presence of virus was examined in lung homogenates. SARS-CoV-2 RNA was measured by RT-qPCR using *RdRP* gene-specific primers ^36^ (S. Table 2), and infectious virus levels were titered on Vero E6 cells. SARS-CoV-2 viral RNA was detected in the lungs of all immunized mice after challenge, with the ATU2 group showing an average 2log_10_ reduction and the ATU3 group a 1log_10_ reduction compared to the empty MV control group (Fig. 6e). However, no infectious virus was detected in the lungs of the ATU2 group and all but one of the ATU3 group (Fig. 6f). These results demonstrate that, although viral replication occurred at low levels, infectivity of the inoculated and progeny virus was efficiently neutralized.

### Partial protection from intranasal challenge after a single immunization

We next determined whether a single immunization could protect IFNAR ^−/−^ mice from challenge with the MACo3 virus (Fig. 7a). Immunized animals were examined for immune responses on days 28 and 48 post-immunization, prior to challenge. All animals exhibited MV-and S-specific antibodies (Fig. 7b, c). Th1-associated IgG responses to the S antigen as well as SARS-CoV-2 NAbs were present before challenge, although at lower levels than after two immunizations (Fig. 7d, e). Mice were then challenged intranasally and lung samples collected 3 days after challenge. Although no difference was observed in viral RNA levels between the test and control groups (Fig. 7f), half of the animals immunized with MV-ATU2-SF-2P-dER were negative for infectious virus in the lungs (Fig. 7g), indicating partial protection. In contrast, animals immunized with MV-ATU3-SF-2P-dER were not protected.

**Fig 7.**
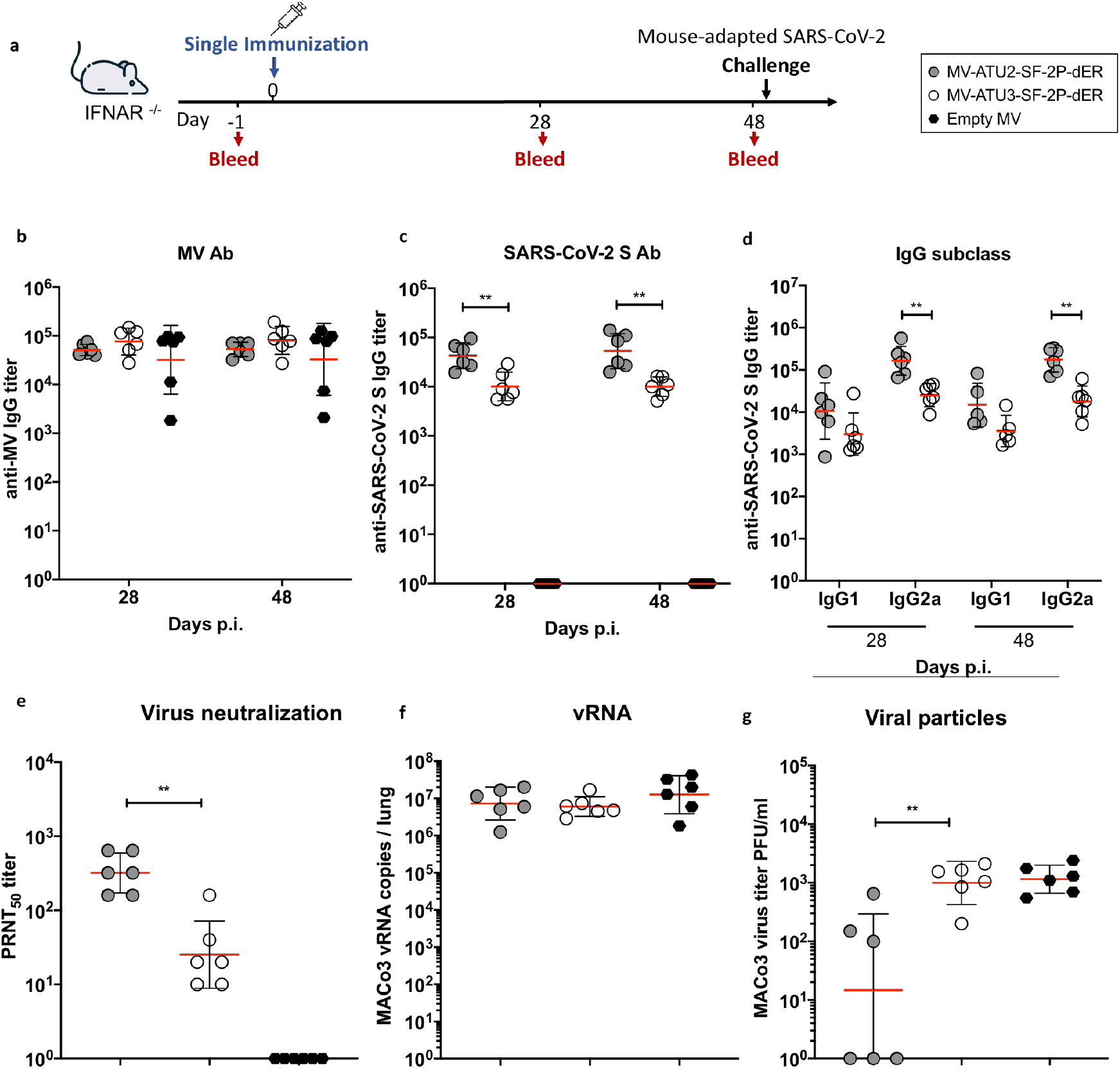
Immune responses and protection after a single immunization. **a)** Immunization and challenge schedule for IFNAR ^−/−^ mice (*n*=6). Animals were immunized interperitoneally on day 0. Sera were collected at days 24 and 48. Animals were challenged on day 48 by intranasal inoculation of mouse-adapted SARS-CoV-2 virus (MACo3) at 1.5 x 10 ^5^ PFU. Sera were assessed for levels of specific antibodies to **b)** MV and **c)** S-SARS-CoV-2 protein. **d)** Neutralizing antibody responses against SARS-CoV-2 virus, expressed as 50% plaque reduction neutralization test (PRNT_50_) titers. **e)** SARS-CoV-2 viral RNA copies detected by RT-qPCR in homogenized lungs of challenged animals, calculated as copies/lung. **f)** Titer of infectious viral particles recovered from the homogenized lung of the immunized animals expressed as PFU/lung. Data are represented as geometric means with line and error bars indicating geometric SD. Statistical significance for antibody responses (top panels) was determined by two-way ANOVA adjusted for multiple comparisons. The rest of the data (bottom panels) was analyzed by the Mann-Whitney *U*-test. Asterisks (*) indicate significant mean differences (* *p*<0.05; ** *p*<0.01, and **** *p*<0.001).

## Discussion

Here we report the development and testing of MV-based COVID-19 vaccine candidates targeting the SARS-CoV-2 S protein. Similar to other vaccine platforms, the full-length prefusion-stabilized S was the most immunogenic, eliciting the strongest humoral and cellular responses. We successfully rescued genectically stable rMV expressing high levels of S antigen from a strong early promoter of measles virus genome. This is in contrast to recently reported MV-based COVID-19 vaccine candidates ^37^. Our lead candidate MV-ATU2-SF-2P-dER exibited a high viral titer (10 ^6^ TCID_50_) comparable to wild-type Schwarz MV strain and was genetically stable up to 10 passages while the previously reported rMV expressing the native S in the same promoter showed impaired growth ^37^. One explaination could be that the 2P mutation helps a stable S protein expression therefore allowing rMV to grow better. Furthermore, MV-ATU2-SF-2P-dER candidate elicited high levels of neutralizing antibodies to SARS-CoV-2 and strong Th1-oriented T-cell reponses. Prime-boost immunization afforded protection from intranasal challenge with a mouse-adapted SARS-CoV-2 virus. Moreover, NAb titers persisted months after the immunization - such long-lasting immunity is a hallmark of replicating vector vaccines ^38^. T-cell responses, essential to controlling and reducing viral load and viral spread ^39^, were induced within seven days after a single immunization. Notably, dominance of Th1 responses suggests that these vaccine candidates are less likely to induce immunopathology due to vaccine-induced disease enhancement as previously reported for SARS-CoV-1 and MERS-CoV vaccine studies ^40, 41, 42, 43, 44^. In addition, after a single immunization, our lead candidate MV-ATU2-SF-2P-dER also provided sufficient immune protection according to WHO recommendations for a COVID-19 vaccine primary efficacy of at least 50% ^45^. This suggests that our lead vaccine candidate can protect against both SARS-CoV-2 infection and disease.

To explore the possibility of generating a broad-spectrum vaccine, we also tested a vaccine candidate expressing only the S2 subunit, which is highly conserved among SARS-CoV-1 and SARS-CoV-2 viruses. The S2 subunit has been shown to harbor immunodominant and neutralizing epitopes ^16, 46, 47^. In this report, while S2 in the MV context induced high S-specific antibody titers, these antibodies could not neutralize the SARS-CoV-2 virus. In terms of cellular responses, S2 did not induce S-specific CD4 ^+^, CD8 ^+^ or IL-5 ^+^ T-cell responses. These observations suggest that the S2 subunit alone is insufficient for inducing immune protection. Given the high titers of non-neutralizing antibodies, it would be interesting to characterize their role in immune responses to SARS-CoV-2 infection and investigate whether they may contribute to immunopathology.

Our results also yielded interesting differences in the immunogenicity of rMV vaccines expressing the target antigen from ATU2 versus ATU3. S antigen is expressed at higher levels from ATU2, and this correlated with higher humoral and cellular responses. Reducing the immunization dose of the ATU2 candidate to 1×10 ^4^ TCID_50_ still induced higher NAb titers than the ATU3 vaccine at 1×10 ^5^ TCID_50_ (S. Fig.8). Additionally, T-cell responses to the MV vector was also lower with ATU2 constructs. These observations suggest that higher antigen expression could be reducing virus replication *in vivo*, resulting in lower cellular responses to the vector itself. While MV vaccines have been shown to be effective despite pre-existing immunity to the vector, this more desirable balance in the immunogenicity of antigen and vector likely contributes to greater vaccine efficacy of the ATU2 construct. Nevertheless, as an rMV vehicle for future vaccines, the ATU3 concept is still useful for expressing antigens that are unstable, toxic, or otherwise difficult to express.

Among the over 200 COVID vaccine candidates currently in development ^48^ the most advanced platforms include human-and chimpanzee-based adenovirus vectors (Janssen, CanSino Biologics, Gamaleya and AstraZeneca), mRNA (Pfizer and Moderna), and inactivated vaccines (SinoVac and SinoPharm). While they already show excellent efficacy results, there are several advantages to MV-vectored vaccines that argue for a strong push to their development.

First, as a replicating vector, MV-based vaccines can be administered at doses as low as 1×10 ^5^ TCID_50_, compared to adenovirus-vectored vaccines that are generally administered at 10 ^10^– 10 ^11^ viral particles. Such high doses can run the risk of strong complement activation ^49^, and the preliminary report from phase I/II trials of AstraZeneca’s ChAdOx1 indicated generally higher, though acceptable, levels of adverse effects even with prophylactic paracetamol administration ^50^. The lower doses relative to production titers render the vaccine more cost-effective for large-scale manufacturing as well. Additionally, replicating vectors are known to provide long-lasting immunity ^51, 52, 53^. Immune responses to our MV-ATU2-SF-2P-dER candidate lasted up to 4 months after prime-boost immunization with no sign of decline, reflecting results seen for the MV-based CHIKV vaccine ^19^. By efficiently stimulating long-lasting memory B-and T-cells, MV induces both humoral and cellular immune responses and provides life-long immunity. MV-CHIKV has an efficacy rate of approximately 90% after one administration and 100% after two administrations.

Second, compared to inactivated and mRNA vaccines, formulation and distribution of MV-vectored vaccines is much simpler. The vector has been shown to naturally activate innate immunity and possess self-adjuvanting properties ^54^, and vaccination with live attenuated rMV therefore does not require adjuvants. Common adjuvants such as alum have been characterized to stimulate Th2-biased responses ^35^, which could increase the possibility of immnunopathology. Furthermore, rMV vaccines can be distributed with the existing cold-chain infrastructure, while mRNA vaccines, which require ultra-cold temperatures will prove to be much more challenging.

Third, while adenovirus-vectored vaccines have to overcome strong pre-existing immunity to the vector, previous clinical studies demonstrated that immune responses to MV-CHIKV were not dampened even when all volunteers were pre-immune to measles ^19, 20^. MV-CHIKV has shown excellent safety and tolerability in both phase I and phase II clinical trials. The success of MV-CHIKV thus far reaffirms that the MV vector is an excellent platform for vaccine development.

Interestingly, the MV vaccine is also known for its non-specific effects (NSEs) in reducing children’s mortality due to other viral infections ^55, 56, 57^. This concept is being tested to see if there is a reduction in the severity of COVID-19 symptoms in a clinical trial of healthcare workers in Egypt (ClinicalTrials.gov. Identifier: NCT04357028. Measles Vaccine in HCW (MV-COVID19) 2020, https://clinicaltrials.gov/ct2/show/NCT04357028).

Given the large percentage of infections resulting in mild or no symptoms ^58^, convincing people to accept vaccination in sufficient numbers worldwide to control the COVID-19 pandemic will be no easy task. Adenovirus vectors and mRNA are both new platforms that have yet to be approved as general-use vaccines, and their accelerated development timelines may become a barrier to acceptance. The long safety track record of the measles vaccine along with its familiarity due to global deployment, however, can help overcome such vaccination barriers. Taken together with our highly promising pre-clinical data, our MV-based SARS-CoV-2 vaccine candidate is an important platform for addressing the COVID-19 pandemic.

## Methods

### Cells and viruses

Human embryonic kidney cells (HEK) 293T (ATCC CRL-3216), HEK293T7-NP helper cells (stably expressing MV-N and MV-P genes), African green monkey kidney cells (Vero) and Vero C1008 clone E6 (ATCC CRL-1586) were maintained at 37°C, 5% CO_2_ in Dulbecco’s modified Eagle medium (DMEM) (Thermo Fisher) supplemented with 5% (for Vero cells) or 10% (for HEK293T cells) heat-inactivated fetal bovine serum (FBS) (Corning), 100 units/ml of penicillin-streptomycin and 100 ug/ml of streptomycin. The SARS-CoV-2 BetaCoV/France/IDF0372/2020 strain was supplied by the National Reference Centre for Respiratory Viruses hosted by Institut Pasteur (Paris, France) and headed by Pr. Sylvie van der Werf. The human sample from which strain BetaCoV/France/IDF0372/2020 was isolated has been provided by Dr. X. Lescure and Pr. Y. Yazdanpanah from the Bichat Hospital, Paris, France. The Mouse-adapated SARS-CoV-2 (MACo-3) has been described elsewhere (Montagutelli et al, in preparation).

### Construction of pTM-MVSchwarz expressing modified SARS-CoV-2 S protein constructs

The SARS-CoV-2 spike (S) gene based on the sequence published by Zhou, Yang ^4^ was codon-optimized for expression in mammalian cells. Primers introducing restriction sites BsiWI and BssHII to the S 5’ and 3’ ends, respectively, were used to amplify nucleotides 1-3799 to generate full-length S (SF) with a deletion of its 11 C-terminal amino acids (SF-dER) (Fig. 1) for cloning into pCDNA 5.1. To generate S2 constructs, primers were designed for inverted PCR with BsmBI restriction sites and 4-nucleotide overlaps at the C-terminus of the native S signal peptide and S2 immediately adjacent to the furin cleavage site (Supplementary Table 2). The amplification product, comprising the S2 region, the pCDNA backbone, and the S signal peptide, was digested with BsmBI (NEB) and self-ligated to generate S2-dER. To maintain the conformation of S in the prefusion state, two mutations were introduced at the hinge of HR1, K986P and V987P (2P mutation) (Fig.1). Primers introducing the mutations were designed with mutated overlapping nucleotides and BsmBI sites (S. Table 2). The SF-dER and S2-dER constructs were amplified, digested and self-ligated to create the prefusion-stabilized SF-2P-dER and S2-2P-dER constructs. The S constructs in the pCDNA background were transfected into Vero cells using FugeneHD. Transfected cells were observed at 24-and 48-hour post-transfection for fusogenic phenotypes.

All S genes were subsequently cloned into pTM-MVSchwarz encoding infectious MV cDNA corresponding to the anti-genome of the MV Schwarz vaccine strain. All the inserted genes were modified at the stop codon to ensure that the total number of nucleotides is a multiple of six ^59^.

### Virus rescue, propagation and titration

Rescue of recombinant MV viruses was performed using a helper-cell-based system as decribed previously ^32^. Briefly, helper HEK293T7-NP cells were individually transfected with 5 µg of pTM-MVSchwartz-based SARS-CoV-2 S plasmids and 0.02 µg of pEMC-La, plasmid expressing the MV polymerase (L) gene ^60^. After overnight incubation at 37°C, the transfection medium was replaced with fresh DMEM medium. Heat shock was applied for 3 h at 42°C before transfected cells were returned to the 37°C incubator. After two days, transfected cells were transferred to 100-mm dishes seeded with monolayers of Vero cells. Syncytia that appeared after 2-3 days of co-culture were singly picked and transferred onto Vero cells seeded in 6-well plates. Infected cells were trypsinized and expanded in 75-cm ^2^ and then 150-cm ^2^ flasks, in DMEM with 5% FBS. When syncytia reached 80%–90% coverage (or when the maximum cytopathic effect (CPE) was observed, usually within 36-48 hours post infection), cells were scraped into a small volume of OptiMEM (Thermo Fisher). Cells were lysed by a single freeze-thaw cycle and cell lysates clarified by low-speed centrifugation. The infectious supernatant was then collected and stored at − 80°C.

Titers of the rMVs were determined on Vero cells seeded in 96-well plates at 7500 cells/well, and infected with serial ten-fold dilutions of virus in DMEM with 5% FBS. After incubation for 7 days, cells were stained with crystal violet, and TCID_50_ values were calculated using the Karber method ^61^. Titers of SARS-CoV-2 and MACo3 were assessed on Vero-E6 in a similar plaque assay. The number of plaques were read 3 days post-infection.

Virus growth kinetics of rMVs was studied on monolayers of Vero cells in 6-well plates. Cells were infected with rMVs at an MOI of 0.1. One plate was used per rMV construct. At various time points post-infection, infected cells were scraped into 1 ml OptiMEM, lysed by freeze-thaw, clarified by centrifugation, and titered as described above.

To assess the stability of S antigen expression by recombinant viruses, Vero cells were repeatedly infected for ten passages. Virus was collected by a freeze-thaw cycle after 1, 5 and 10 passages (P1, P5, P10) and used to infect Vero cells in 6 well-plates in duplicate. Cell lyzates were then assessed for S mRNA and protein levels using RT-PCR, western blotting, and NGS respectively.

## RT-PCR

To verify S expression from the rMV constructs, total RNA were extracted from infected Vero cells using the RNeasy Mini Kit (Qiagen). The cDNA synthesis and PCR steps were performed using the RNA LA PCR kit (Takara Bio) with primers (S. Table 2) targeting ATU2 and ATU3, according to the manufacturer’s instructions. RT-PCR products were verified by Sanger sequencing (Eurofins Genomics) using the primers indicated in S. Table 2.

### Western blot analysis

Vero cells in 6-well plates were infected with various rMVs at an MOI of 0.1. At 36–48 h post-infection (80% syncytia), infected cells were lysed in RIPA lysis buffer (Thermo Fisher). Samples were briefly centrifuged and subjected to SDS-PAGE using the NuPAGE-pre-cast 4–12% gradient gel with NuPAGE-MOPs running buffer (Invitrogen). After transfer to a nitrocellulose membrane (GE Healthcare) and blocking with Tris-buffered saline (TBS) buffer with 0.1% Tween, 5% milk, the membrane was subsequently probed with a rabbit polyclonal anti-SARS-CoV S antibody recognizing the conserved 1124 aa–1140 aa epitope (ABIN199984, Antibody Online, 1:2000 dilution) followed by a horse-radish peroxidase (HRP)-conjugated swine anti-rabbit IgG antibody (P0399, Dako, 1:3000 dilution). Bands were visualized using SuperSignal West pico Plus chemiluminescent HRP substrate (Thermo Fisher). For loading control, membranes were stripped with 5% NaOH for 5 mins then blocked. Membranes were then re-probed with a mouse monoclonal anti-MV-N antibody (ab9397, Abcam, 1:20000 dilution) followed by an HRP-conjugated anti-mouse IgG (NA931V, GE Healthcare, 1:10000 dilution).

### Next-Generation Sequencing

Extracted RNA was treated with Turbo DNase (Ambion) followed by purification using SPRI beads (Agencourt RNA clean XP, Beckman Coulter). We used a ribosomal RNA (rRNA) depletion approach based on RNAse H and targeting human rRNA. The RNA from the selective depletion was used for cDNA synthesis using SuperScript IV (Invitrogen) and random primers, followed by second-strand synthesis. Libraries were prepared using a Nextera XT kit and sequenced on an Illumina NextSeq500 (2×150 cycles) at the Mutualized Platform for Microbiology hosted at Institut Pasteur. Raw reads were trimmed using Trimmomatic v0.39 ^62^ to remove adaptors and low-quality reads. We assembled reads using metaspades v3.14.0 ^63^ with default parameters. Scaffolds were queried against the NCBI non-redundant protein database (13) using DIAMOND v2.0.4 (14). No other viruses were detected. The recombinant MV genomes identified were verified and corrected by iterative mapping using CLC Assembly Cell v5.1.0 (QIAGEN). Aligned reads were manually inspected using Geneious prime v2020.1.2 (2020) (https://www.geneious.com/), and consensus sequences were generated using a minimum of 5X read-depth coverage to make a base call. Minor variants frequencies were estimated using Ivar _64_.

### Immunofluorescence assay

Vero cells were infected with various rMVs at an MOI of 0.1. At 24–36h post-infection, cells were fixed with 4% paraformaldehyde, blocked with 2% goat serum overnight and then treated with or without 0.1% saponin A (Sigma). Fixed cells were then probed with a mouse monoclonal anti-SARS-CoV S antibody (ab273433, Abcam, 1:300 dilution) as the primary antibody. An Alexa Fluor 488-conjugated goat anti-rabbit IgG (A-11008, Thermo Fisher) was used as the secondary antibody. Staining with anti-MV-N followed by Cy3-conjugated goat anti-rabbit (A10520, Jackson ImmunoResearch, 1:1000 dilution) was used to detect MV in the same infected cells. Nuclei were stained with DAPI. Images were collected using an inverted Leica DM IRB fluorescence microscope with a 20x objective.

### Flow cytometry

pcDNA5.1 expression vectors encoding prefusion-stabilized or native conformation full-length S and S2 subunit antigens were used to transfect HEK293T cells using the JetPrime transfection kit (PolyPlus) according to the manufacturer’s instructions. Forty-eight hours post-transfection, cells were stained for indirect immunofluorescence with 10 μg/ml of rabbit polyclonal anti-S antibody targeting S2 (ABIN199984) followed by Alexa Fluor 488-conjugated goat anti-rabbit IgG (A-11008). Propidium iodide was used to exclude dead cells by gating. Stained cells were acquired on the Attune NxT flow cytometer (Invitrogen) and data were analyzed using FlowJo v10.7 software (FlowJo LLC).

### Mice immunizations and challenge

All experiments were approved by the Office of Laboratory Animal Care at the Institut Pasteur and conducted in accordance with its guidelines. Groups of 6 to 8-week-old mice deficient for type-I IFN receptor (IFNAR ^−/−^) were intraperitoneally injected with 10 ^5^ TCID_50_ rMV, namely SF-2P-dER or S2-2P-dER in ATU2 or ATU3, or the control empty MV Schwarz. To study humoral responses, two immunizations were administered at a four-week interval. Sera were collected before the first immunization (day-1) and then before (day 28) and after (day 42) after the second immunization. All serum samples were heat-inactivated for 30 min at 56 °C. To assess protection, mice that received either one or two immunizations were challenged with an intransal inoculation of 1.5 x 10 ^5^ PFU mouse-adapted SARS-CoV-2 virus (MACo3). Three days after challenge, mice were sacrificed and lung samples collected. The presence of MACo3 virus in the lung was detected by determining viral growth, PFU of infectious viral particles, and measuring vRNA using Luna Universal Probe One-Step RT-qPCR kit following the manufacturer protocol (E3006). The primers and probes used correspond to the nCoV_IP4 panel (S.Table2) as described on the WHO website ^36^.

## ELISA

Edmonston strain-derived MV antigens (Jena Bioscience) or recombinant S protein encompassing amino acid residues 16 to 1213 with R683A and R685A mutations (ABIN6952426, Antibodies Online) were coated on NUNC MAXISORP 96-well immuno-plates (Thermo Fisher) at 1µg/ml in 1x phosphate-buffered saline (PBS). Coated plates were incubated overnight at 4°C, washed 3 times with washing buffer (PBS, 0.05% Tween), and further blocked for 1h at 37°C with blocking buffer (PBS, 0.05% Tween, 5% milk). Serum samples from immunized mice were serially diluted in the binding buffer (PBS, 0.05% Tween, 2.5% milk) and incubated on plates for 1h at 37°C. After washing steps, an HRP-conjugated goat anti-mouse IgG (H+L) antibody (Jackson ImmunoResearch, 115-035-146, 1:5000 dilution) was added for 1h at 37°C. Antibody binding was detected by addition of the TMB substrate (Eurobio) and the reaction was stopped with 100 µl of 30% H_2_SO_4_. The optical densities were recorded at 450 and 620 nm wavelengths using the EnSpire 2300 Multilabel Plate Reader (Perkin Elmer). Endpoint titers for each individual serum sample were calculated as the reciprocal of the last dilution giving twice the absorbance of the negative control sera. Isotype determination of the antibody responses was performed using HRP-conjugated isotype-specific (IgG1 or IgG2a) goat anti-mouse antibodies (AB97240 and AB97245, Abcam, 1:5000).

### Plaque reduction neutralization test

Two-fold serial dilutions of heat-inactivated serum samples were incubated at 37 °C for 1 h with 50 PFU of SARS-CoV-2 virus in DMEM medium without FBS and added to a monolayer of Vero E6 cells seeded in 24-well plates. Virus was allowed to adsorb for 2 h at 37 °C. The supernatant was removed and the cells were overlaid with 1 ml of plaque assay overlay media (DMEM supplemented with 5% FBS and 1.5% carboxymethylcellulose). The plates were incubated at 37 °C with 5% CO_2_ for 3 days. Viruses were inactivated and cells were fixed and stained with a 30% crystal violet solution containing 20% ethanol and 10% formaldehyde (all from Sigma). Serum neutralization titer was counted on the dilution that reduced SARS-CoV-2 plaques by 50% (PRNT_50_).

## ELISPOT

Splenocytes from immunized mice were isolated and red blood cells lysed using Hybri-Max Red Blood Cell Lysing Buffer (Sigma). The splenocytes were tested for their capacity to secrete IFN-γ upon specific stimulation. Multiscreen-HA 96-well plates (Millipore) were coated overnight at 4°C with 100 µl per well of 10µg/ml of anti-mouse IFN-γ (551216, BD Biosciences) in PBS before washing and blocking for 2h at 37°C with 200 µl complete MEM-α (MEM-α (Thermo Fisher) supplemented with 10% FBS, 1x non-essential amino acids, 1mM sodium pyruvate, 2mM L-glutamine, 10mM HEPES, 1% penicillin-streptomycin, and 50 μM β-mercaptoethanol). The medium was replaced with 100µl of cell suspension containing 1×10 ^5^ splenocytes in each well in triplicate and 100µl of stimulating agent in complete MEM-α supplemented with 10 U/ml of mouse IL-2 (Roche). Stimulating agents used were 2.5 µg/ml concanavalin A (Sigma Aldrich) for positive controls, complete MEM-α for negative controls, MV Schwarz virus at an MOI of 1, or a SARS-CoV-2 S peptide pool (S. Table 1) at 2 µg/ml per peptide. After incubation for 40h at 37°C, 5% CO_2_, plates were washed once with PBS, then three times with washing buffer (PBS, 0.05% Tween). A biotinylated anti-mouse IFN-γ antibody (554410, BD Biosciences) at 1µg/ml in the washing buffer was added and plates were incubated for 120min at room temperature. After extensive washing, 100 µl of streptavidin–alkaline phosphatase conjugate (Roche) was added at a dilution of 1:1000 and plates were further incubated for 1h at room temperature. Wells were washed twice with the washing buffer and followed by a wash with PBS buffer without Tween. Spots were developed with BCIP/NBT (Sigma) and counted on a CTL ImmunoSpot® ELISPOT reader.

### Intracellular cytokine staining

Splenocytes of vaccinated mice were extracted as previously described. Two million splenocytes per mouse per well were incubated in 200 μL of complete MEM-α medium (Thermo Fisher). BD Golgi Stop (554724, BD Biosciences) was added to the culture medium according to the manufacturer’s instructions. Splenocytes were stimulated with a peptide pool covering the predicted CD4 and CD8 T-cell epitopes of the SARS-CoV-2 S protein (S. Table 2) at a final concentration of 2 µg/ml per peptide. PMA/Ionomycin Cell Stimulation Cocktail (eBioscience) was used as a stimulation for positive controls, and medium alone was used for negative controls. Splenocytes were stimulated for 4 h at 37 °C. Stimulated cells were incubated with Mouse BD Fc Block (553141, BD Biosciences), and stained with Live/Dead Fixable Aqua Viability Dye (ThermoFisher) to exclude dead cells by gating. Subsequently, cells were stained with CD3e PE (clone 145-2C11, 12-0031-83, eBioscience), CD4 PerCP-eFluor710 (clone RM4-5, 46-0042-82) and CD8 Alexa Fluor 488 (clone 53-6.7, 53-0081-82) antibodies from Invitrogen. Cells were fixed and permeabilized with BD Fixation/Permeabilization kit (BD Biosciences) and stained with IFN-γ APC/Fire750, (clone XMG1.2, 505860, BioLegend), TNF-α BV421 (clone MP6-XT22, 563387, BD Horizon), IL-5 APC (clone TRFK5, 505860, BD Biosciences), IL-13 eFluor 660 (clone eBio13A, 50-7133-82, eBioscience), CD62L APC-Cy7 (clone MEL-14, 104427, BioLegend), and CD44 BV786 (clone IM7, 563736, BD Pharmingen) antibodies. Samples were acquired using the Attune NxT flow cytometer (Invitrogen) and data were analyzed using FlowJo v10.7 software (FlowJo LLC).

### Statistical information

Statistical analyses were performed using GraphPad Prism v.8.0.2. Results were considered significant if p < 0.05. The lines in all graphs represent the geometric mean with error bars indicating geometric SD. Statistical analyses of antibody responses, ELISA and PRNT_50_, were done using two-way ANOVA adjusted for multiple comparisons. The two-tailed nonparametric Mann-Whitney’s *U* test was applied to compare differences between two groups.

### Data Availability

The authors declare that the data supporting the findings of this study are available within this article and its supplementary information files or available from the corresponding author upon reasonable request. The source data files are provided with this paper.

## Acknowledgments

We thank Vincent Enouf and Sylvie Van Der Werf from the RNA Viruses Molecular Genetics Unit of Institut Pasteur for helpful discussions and SARS-Cov-2 virus information. We thank Maud Vanpeene and Vincent Enouf from the Mutualized Platform of Microbiology, Pasteur International Bioresources Network for the viral sequencing.

## Author contributions

PNF, AB, AJ, CG and FT conceived the study; PNF, AB, CR, CC, VN, MP, LP, XM, PF, HSM, and ESL performed experiments; PNF, AB, HSM, JDS, ESL and FT analyzed data; PNF and AB performed statistical analysis; PNF, AB, ST and FT wrote the manuscript.

