## supplementary data for "A measles-vectored COVID-19 vaccine induces long-term immunity and protection from SARS-CoV-2 challenge in mice"

### SFig. 1

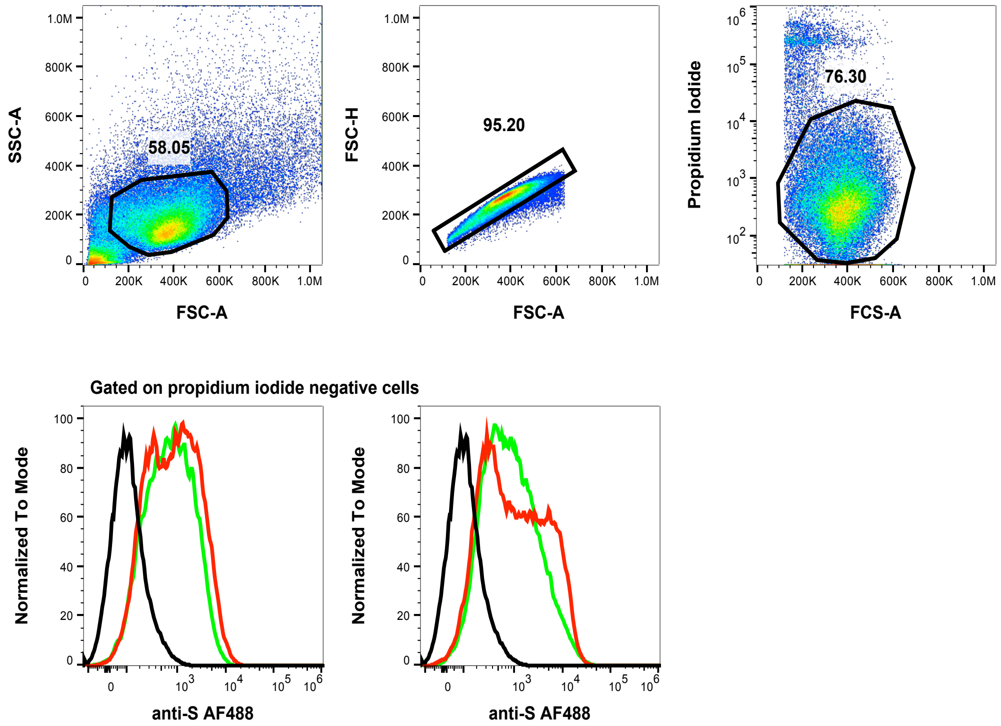

| Sample | MFI |
|---|---|
| Negative control | 178 |
| SF-2P-dER | 1604 |
| SF-dER | 1237 |

| Sample | MFI |
|---|---|
| Negative control | 178 |
| S2-2P-dER | 2687 |
| S2-dER | 1907 |

**Supplementary figure 1.** Expression of SARS-CoV-2 S antigens on the surface of transfected HEK293T cells. Cells transfected with pcDNA expression vectors encoding full-length S or S2 subunit antigens were stained for indirect immunofluorescence with an anti-S antibody followed by Alexa Fluor 488-conjugated goat anti-rabbit IgG. Propidium iodide was used to exclude dead cells by gating (upper dot plots). Histograms show surface expression of full-length S (left histograms) or S2 subunit proteins (right histograms). Native-conformation S antigens (green), prefusion-stabilized S (red), mock-transfected control cells (black histograms) and corresponding mean fluorescence intensities (MFI) are shown.

#### SFig. 2

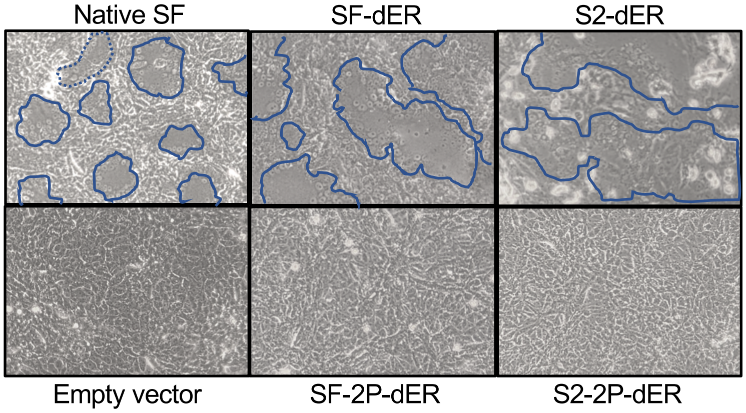

**Supplementary figure 2.** S protein-mediated syncytium formation in transfected Vero cells. Images of Vero cells transfected with pcDNA expression vectors encoding SARS-CoV-2 S proteins were acquired 24 hours post-transfection. Upper images show Vero cells transfected with plasmids encoding native-conformation S antigens, while lower images depict cells transfected with prefusion-stabilized S antigens and non-transfected control Vero cells. Blue lines delineate the borders of syncytia. Native SF indicates native-conformation full-length S protein with an intact CT.

**SFig. 3**

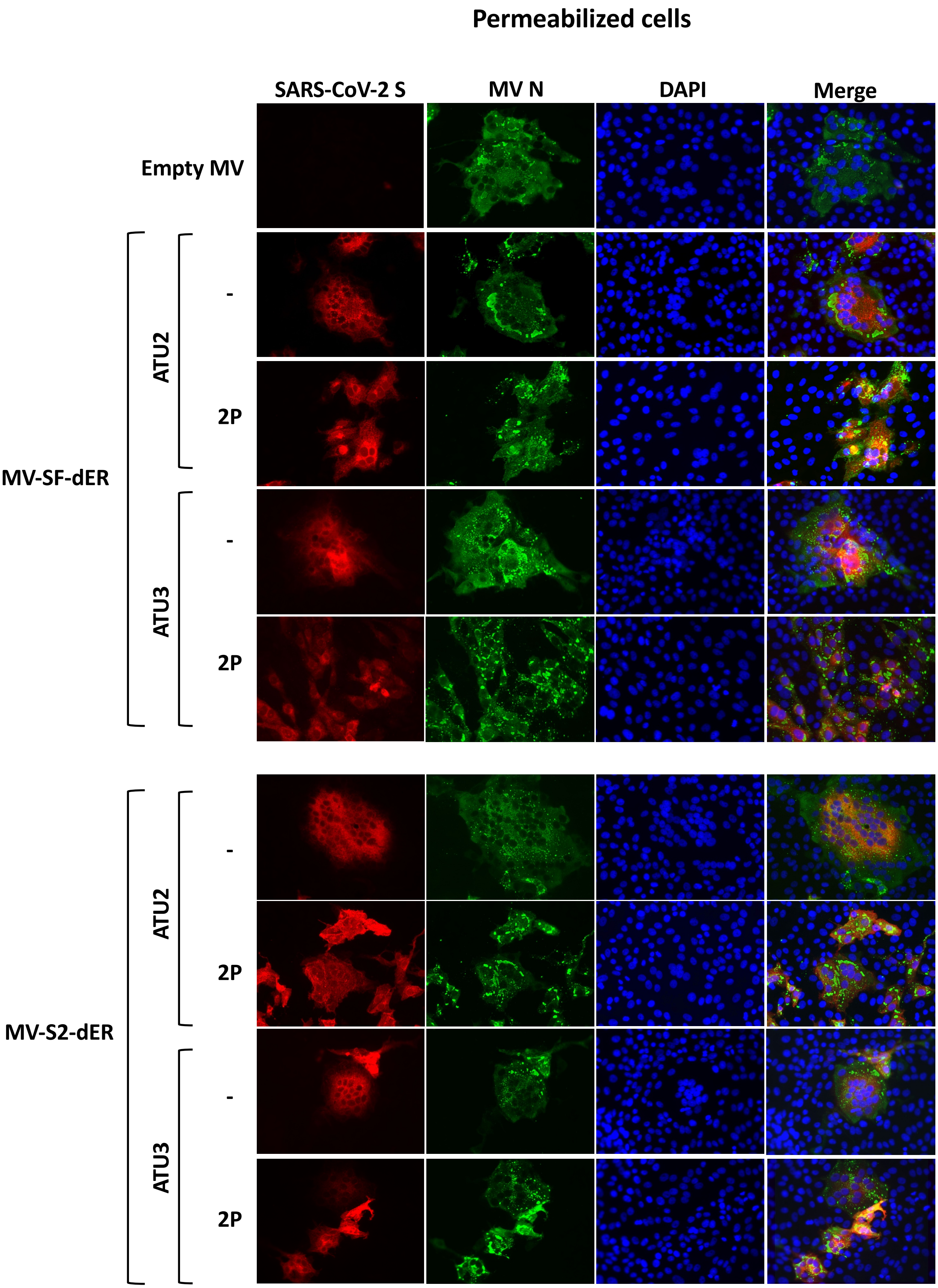

**Supplementary figure 3.** Immunofluorescence analysis of intracellular S protein expression in Vero cells infected with recombinant MV vaccines. Vero cells were infected with rMVs expressing SARS-CoV-2 S proteins or empty MV Schwarz. Twenty-four hours after infection, S protein was detected in saponin-permeabilized cells using a rabbit anti-S antibody followed by Cy3-conjugated goat anti-rabbit IgG (red). MV N protein was visualized using mouse monoclonal anti-N antibody followed by Alexa Fluor 488-conjugated goat anti-mouse IgG (green). Nuclei were stained with DAPI (blue). Images were acquired using a fluorescence microscope.

SFig. 4

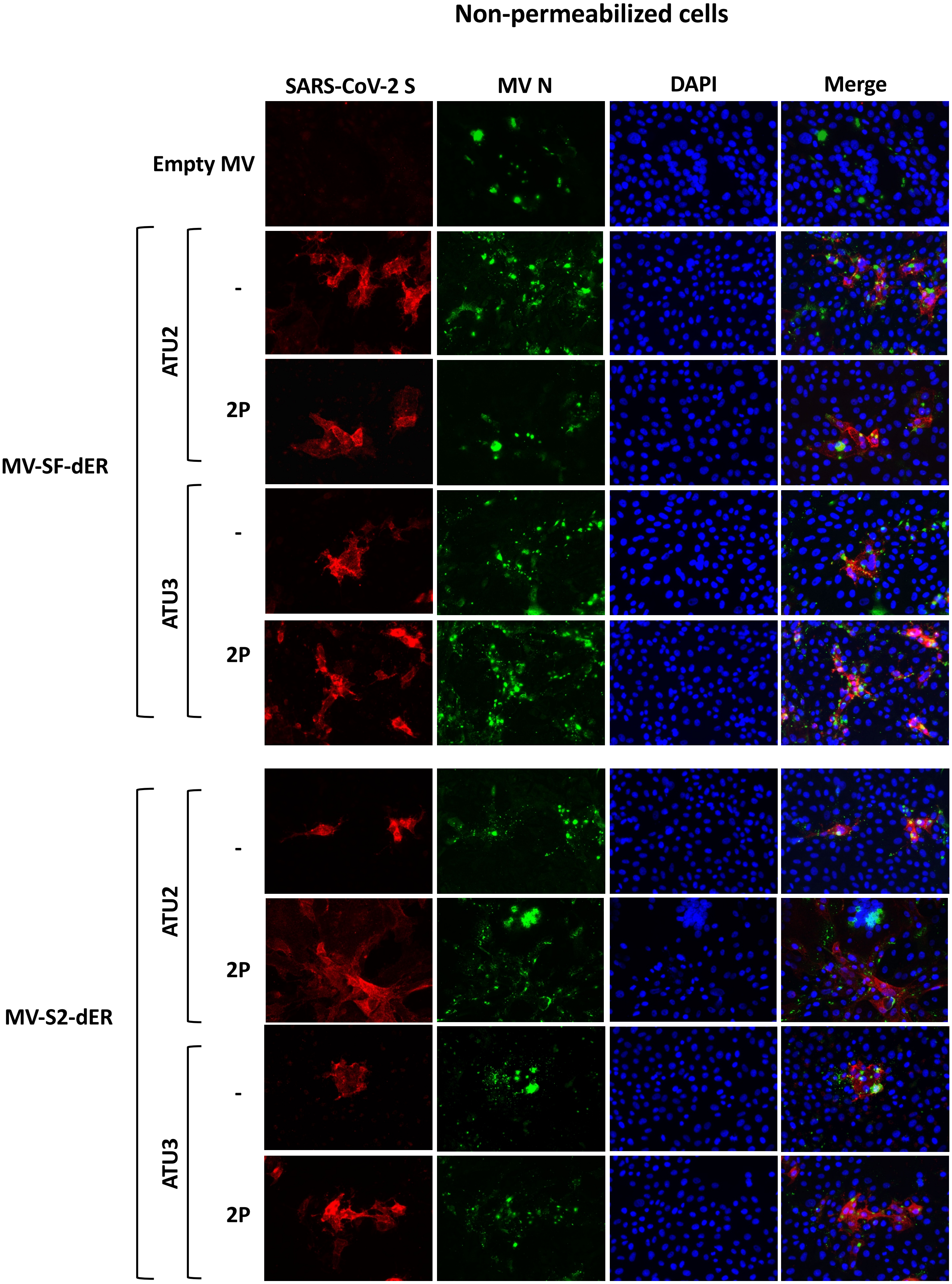

**Supplementary figure 4.** Immunofluorescence analysis of S protein surface expression in Vero cells infected with recombinant MV vaccines. Vero cells were infected with rMVs expressing SARS-CoV-2 S proteins or empty MV Schwarz. Twenty-four hours after infection, S protein was detected on the surface of the non-permeabilized cells using a rabbit anti-S antibody followed by Cy3-conjugated goat anti-rabbit IgG (red). MV N protein was visualized using a mouse monoclonal anti-N antibody followed by Alexa Fluor 488-conjugated goat anti-mouse IgG (green). Nuclei were stained with DAPI (blue). Images were acquired using a fluorescence microscope.

### SFig. 5

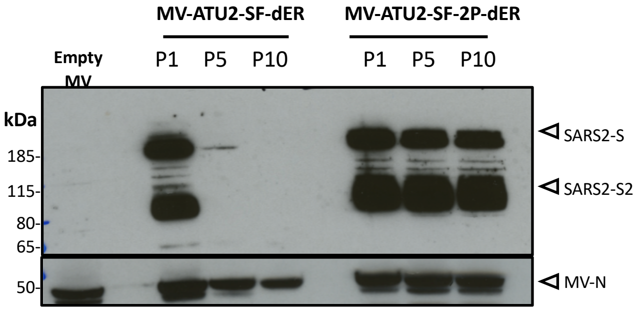

**Supplementary figure 5.** Western blot analysis of S protein expression in Vero cells infected with recombinant MV vaccines from serial passages. MV ATU2 vaccines expressing SF-dER or SF-2P-dER antigens were serially passaged on Vero cells from P1 up to P10 and S protein expression was determined by immunoblotting of P1, P5, and P10 cell lysates. Vero cells infected with empty MV were examined in parallel and served as negative controls.

**SFig. 6**

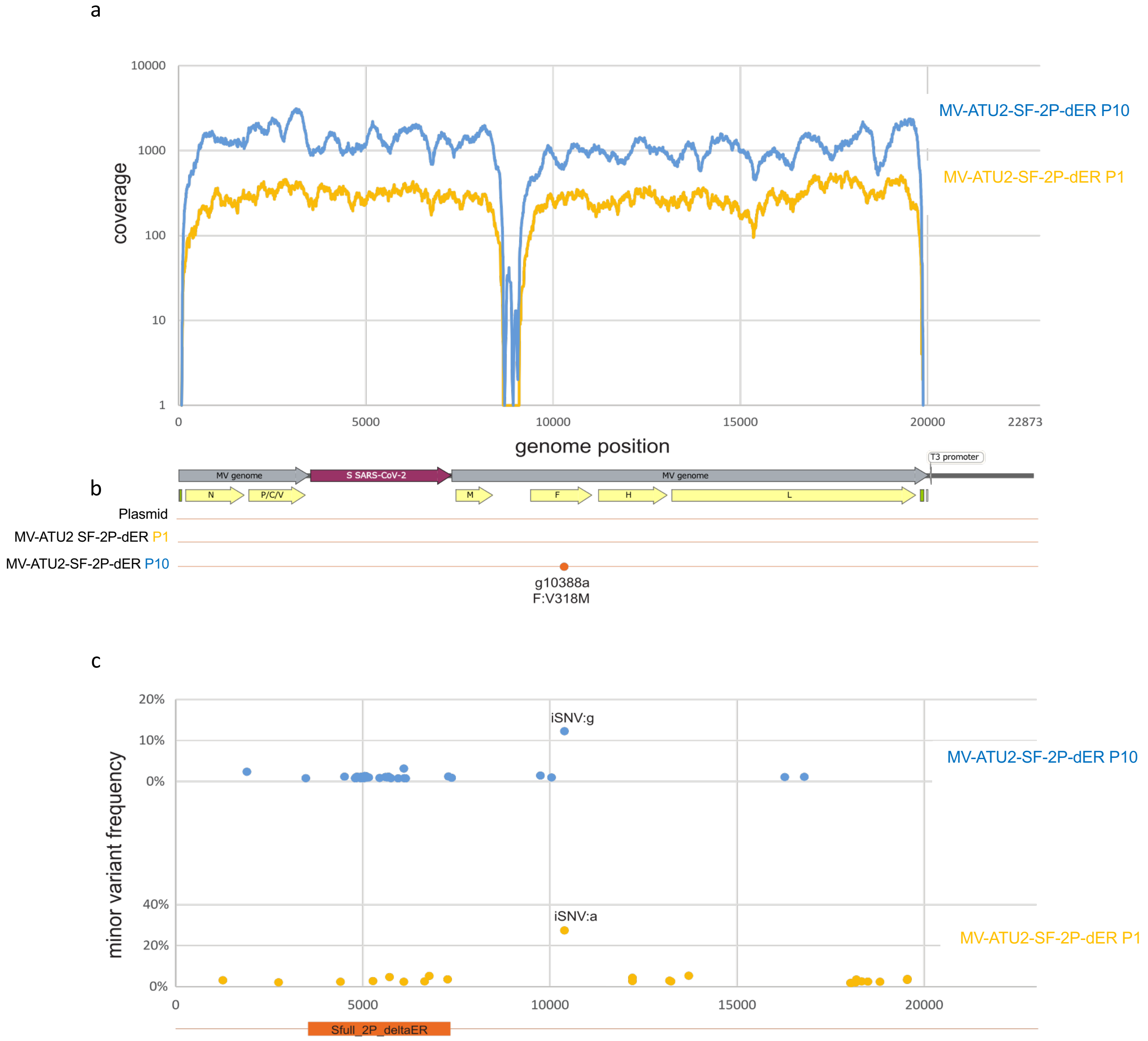

**Supplementary figure 6.** Genome coverage of MV-ATU2-SF-2P-dER from Next-Generation Sequencing depicted in; **a** Schematic of genome coverage to with the genome position with yellow and blue lines represent MV-ATU2-SF-2P-dER genome sequencing of P1 and P10, respectively. **b** Schematic of genome comparison of plasmid sequence to MV-ATU2-SF-2P-dER genome sequencing of P1 and P10. **c** Percent minor variant frequency with yellow and blue dots represent MV-ATU2-SF-2P-dER genome of P1 and P10.

**SFig. 7**

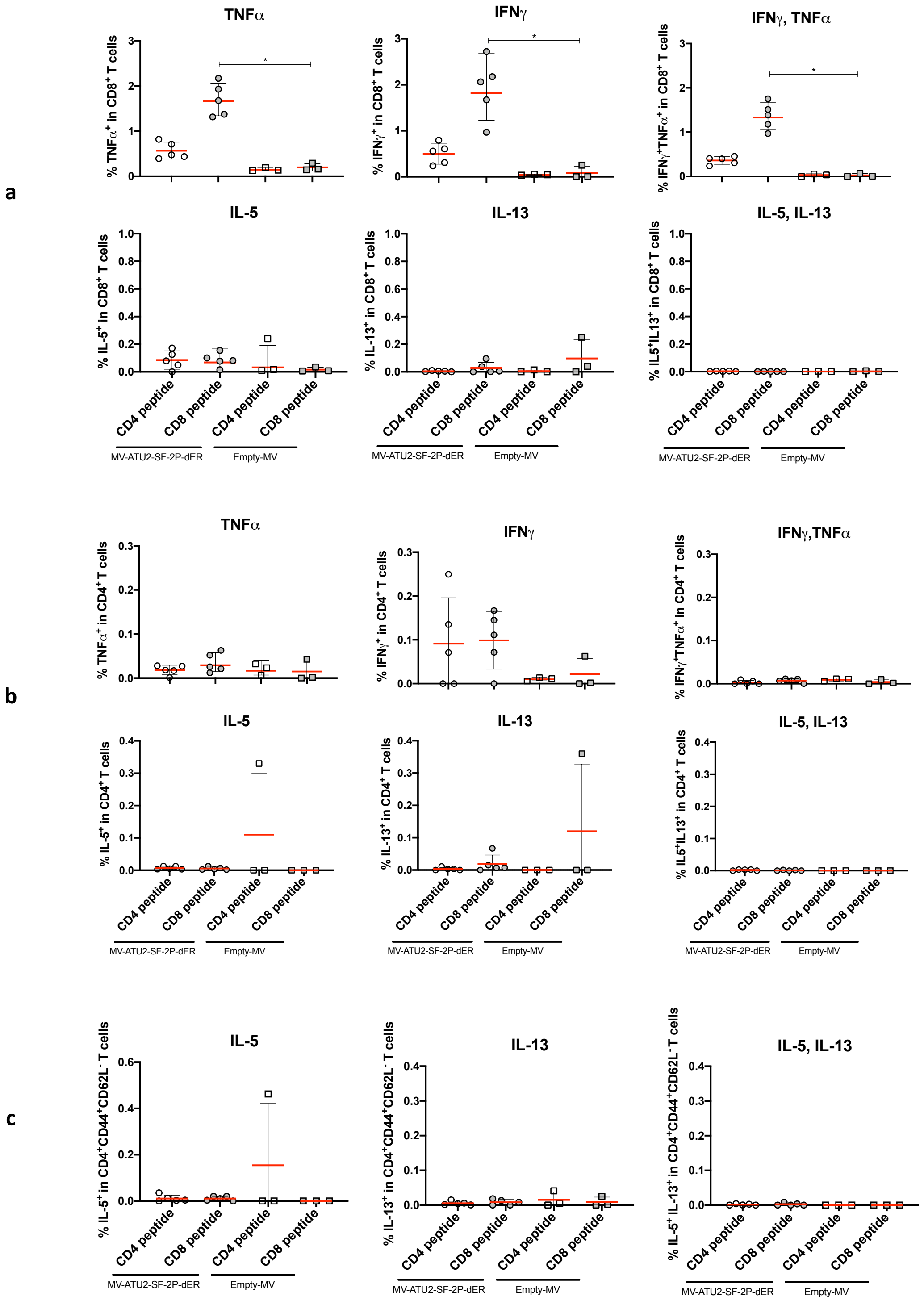

**Supplementary figure 7.** Cytokine expression profile of T cells assessed in IFNAR$^{-/-}$ mice (n=5 or n=3 for a control Empty MV group) immunized intraperitoneally (i.p.) with 1x10$^{5}$ TCID$_{50}$ of MV-ATU2-SF-2P-dER or Empty MV. Splenocytes were stimulated with either S-specific CD4 or CD8 peptide (S. Table1). S-specific **a** CD8$^{+}$ and **b** CD4$^{+}$ T-cells were stained for intracellular IFNγ, TNFα, IL-5 and IL13. **c** S-specific CD4$^{+}$ memory T cells were stained for intracellular IL-5 and IL13. Asterisks (*) indicate significant mean differences (* $p < 0.05$) as determined by Kruskal-Wallis ANOVA with multiple comparisons tests.

**SFig. 8**

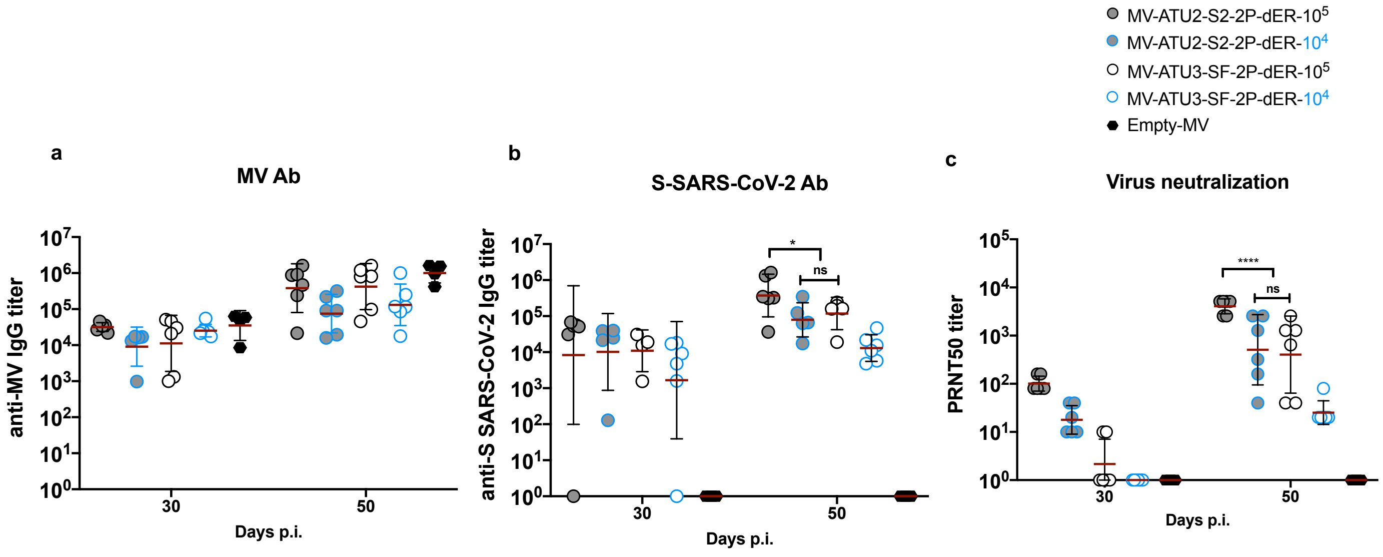

**Supplementary figure 8.** Dose-dependent homologous prime-boost immunization. IFNAR $^{-/-}$ mice (*n*=6 or *n*=4 for the empty MV control) were immunized intraperitoneally with the indicated rMV vaccine candidates at $1\times10^5$ $TCID_{50}$ or $1\times10^4$ $TCID_{50}$ at days 0 and 28. Sera were collected 28 and 50 days after immunization and assessed for specific antibody responses to **a** MV antigen or **b** SARS-CoV-2 S protein. The data show the reciprocal endpoint dilution titers with each data point represents an individual animal. **c** Neutralizing antibody response to SARS-CoV-2 virus expressed as 50% plaque reduction neutralization test ($PRNT_{50}$) titers. Data are represented as geometric means with lines and error bars indicating geometric SD. Statistical significance was determined by a two-way ANOVA adjusted for multiple comparisons. Asterisks (*) indicate significant mean differences (*$p < 0.05$, **$p < 0.01$, and **** $p < 0.001$ ).

### SFig. 9

```
>SF-dER DNA sequence  (3792 bp)
ATGTTCGTCTTTCTGGTATTGCTTCCTCTGGTGAGTTCCCAGTGCGTGAACCTGACGACTCGGACACAGCTGCCGCCTGCATA
CACAAACAGCTTCACACGAGGTGTTTACTATCCGGACAAGGTGTTCCGGTCCTCTGTCTTGCACAGTACTCAGGACCTTTTTC
TGCCATTCTTTAGTAACGTAACATGGTTCCACGCTATCCATGTAAGTGGAACCAATGGCACCAAGCGCTTCGACAATCCAGTC
CTGCCTTTCAACGATGGCGTCTACTTCGCCTCTACCGAGAAATCAAATATTATTAGAGGGTGGATCTTCGGGACCACCCTTGA
TAGCAAGACTCAGAGTCTTCTGATCGTGAACAATGCTACTAACGTCGTCATAAAAGTATGTGAATTTCAGTTCTGCAATGACC
CCTTCCTGGGCGTGTACTATCACAAAAATAACAAGTCCTGGATGGAATCCGAGTTTCGGGTATACAGCAGCGCTAATAACTGT
ACCTTCGAATATGTGAGCCAGCCATTCCTGATGGACCTGGAGGGGAAGCAGGGCAACTTCAAAAACCTGAGGGAATTCGTGTT
CAAGAACATTGACGGATACTTCAAAATCTACAGCAAGCACACTCCAATTAATCTTGTTAGGGACCTTCCTCAGGGGTTCTCCG
CGCTGGAACCCTTGGTGGATCTGCCAATTGGTATTAACATCACGAGGTTTCAAACCCTGCTCGCCCTCCACCGGTCATACCTG
ACCCCAGGTGATAGTAGTAGTGGCTGGACAGCAGGGGCCGCAGCATACTACGTCGGCTACCTCCAGCCCCGCACATTCCTGCT
GAAGTATAATGAAAACGGTACTATTACAGATGCTGTGGACTGCGCACTTGATCCATTGTCCGAGACTAAATGCACCTTGAAGT
CTTTTACCGTTGAGAAGGGAATCTATCAGACCAGTAATTTCAGAGTGCAACCCACCGAGAGCATCGTGCGATTTCCTAATATT
ACAAACCTGTGCCCTTTTGGCGAGGTGTTTAACGCCACTCGCTTCGCGAGCGTGTATGCATGGAACCGGAAAAGGATTAGCAA
CTGCGTGGCTGATTATTCAGTCCTTTATAATAGTGCCTCTTTCAGTACGTTCAAATGCTATGGAGTGTCCCCAACTAAGCTTA
ACGATCTGTGTTTTACCAACGTGTATGCCGACTCTTTCGTCATCAGGGGCGACGAGGTTAGACAAATTGCGCCCGGACAGACA
GGCAAAATTGCTGACTACAATTATAAACTGCCAGATGACTTTACAGGATGCGTGATTGCCTGGAACAGCAACAACCTTGATTC
TAAGGTGGGCGGAAACTACAACTATCTGTACAGGCTGTTCCGCAAATCAAACCTGAAGCCATTCGAAAGGGACATTTCAACTG
AGATCTACCAGGCCGGATCTACACCTTGCAACGGGGTGGAAGGATTCAACTGCTACTTCCCGCTCCAGTCATATGGCTTTCAA
CCTACCAATGGAGTGGGCTACCAACCTTATCGAGTAGTTGTTCTCTCTTTCGAGTTGTTGCATGCCCCAGCCACTGTCTGCGG
ACCAAAGAAATCCACCAATCTGGTGAAAAACAAGTGCGTCAACTTCAACTTCAATGGACTCACCGGTACAGGGGTTCTTACCG
AGTCAAACAAGAAATTCCTTCCGTTTCAGCAGTTTGGACGCGATATCGCCGACACTACGGATGCAGTACGCGATCCTCAGACA
CTTGAGATCCTGGACATCACTCCATGCTCTTTTGGAGGGGTGTCTGTCATCACTCCCGGGACGAATACCAGTAATCAGGTCGC
CGTGCTCTATCAAGATGTCAACTGCACAGAGGTGCCAGTCGCCATTCACGCCGATCAGCTTACCCCTACTTGGCGAGTATATA
GCACAGGGTCCAACGTGTTTCAGACAAGGGCGGGTTGTCTGATCGGCGCTGAGCACGTCAATAATAGCTACGAATGTGACATC
CCTATTGGAGCCGGGATTTGCGCTAGCTATCAGACTCAGACTAACTCACCCCGGCGAGCTAGGTCTGTAGCAAGTCAGAGTAT
CATAGCCTATACTATGAGCCTGGGGGCAGAAAATTCTGTCGCTTATAGTAATAACAGTATTGCAATTCCTACTAACTTCACTA
TTAGCGTGACTACAGAAATTCTGCCAGTGAGCATGACCAAGACCTCCGTCGACTGTACCATGTATATCTGCGGAGACAGCACA
GAATGCAGCAACCTGCTTCTGCAGTACGGTAGCTTTTGTACCCAGCTGAACAGGGCACTGACCGGCATCGCTGTGGAACAGGA
CAAAAATACGCAGGAGGTGTTCGCTCAGGTCAAGCAGATTTATAAGACACCCCCAATAAAAGACTTCGGCGGCTTTAACTTCT
CCCAGATCCTGCCCGATCCGTCAAAACCTTCCAAGAGAAGTTTTATCGAGGATCTGCTGTTTAACAAAGTAACATTGGCTGAT
GCTGGCTTTATTAAACAGTACGGCGATTGTCTGGGAGATATCGCCGCTAGAGATCTGATCTGCGCTCAAAAATTTAATGGCCT
CACTGTATTGCCCCCCCTGCTGACGGATGAAATGATTGCCCAATACACATCAGCCCTGCTGGCTGGGACAATCACTTCCGGCT
GGACGTTCGGTGCTGGAGCTGCTCTTCAAATTCCATTTGCCATGCAAATGGCTTATAGATTCAACGGAATTGGCGTGACCCAA
AACGTGCTGTACGAAAATCAAAAGCTGATTGCTAATCAGTTTAACTCCGCTATCGGTAAGATACAGGACAGCCTGTCTAGTAC
TGCCAGCGCCCTCGGAAAGCTCCAGGATGTGGTGAACCAGAATGCGCAGGCGCTGAACACACTCGTCAAACAGCTTAGCAGCA
ATTTCGGCGCAATTAGCAGTGTACTGAACGACATCTTGTCTCGGCTCGACAAGGTCGAGGCCGAAGTGCAGATTGATCGCCTG
ATCACTGGCCGGCTCCAGTCTCTGCAGACCTACGTAACCCAGCAACTCATACGCGCTGCCGAAATTCGGGCTTCAGCTAACCT
GGCCGCAACAAAGATGTCTGAATGTGTTCTTGGGCAGTCCAAGAGGGTGGACTTCTGCGGGAAAGGATATCACCTGATGTCCT
TCCCTCAGAGCGCCCCTCATGGAGTGGTGTTTCTTCATGTTACCTATGTTCCGGCCCAGGAAAAGAATTTTACCACTGCCCCA
GCCATTTGCCATGATGGTAAAGCACACTTCCCAAGAGAGGGCGTGTTTGTTAGTAACGGCACCCACTGGTTCGTTACCCAGCG
CAATTTTTACGAACCTCAAATCATTACCACTGACAATACATTTGTATCAGGCAATTGCGACGTGGTGATCGGAATCGTAAATA
ATACAGTCTACGATCCCCTGCAGCCAGAGCTTGACAGCTTTAAAGAGGAACTGGACAAATACTTCAAAAATCATACAAGCCCC
GACGTCGACCTGGGAGATATTTCTGGCATCAATGCCTCCGTCGTGAATATCCAAAAAGAGATCGACCGCCTTAATGAGGTTGC
CAAAAACCTCAACGAGTCCCTGATTGATCTGCAGGAGCTGGGGAAGTACGAGCAGTATATCAAATGGCCCTGGTACATCTGGC
TGGGCTTCATTGCCGGGCTCATAGCCATCGTGATGGTGACGATCATGCTGTGTTGCATGACCTCTTGCTGTTCTTGTCTGAAG
GGGTGCTGCTCTTGTGGGAGTTGTTGTAAATTCGATGAGGATGATTCCGAATAATAG
```

**Supplementary figure 9.** DNA sequence of SARS-CoV-2 SF-dER used as the template to construct SARS-CoV-2 vaccine candidates.

### S.Table 1

**Supplementary Table1.** Peptide pools corresponding to the S1 and S2 subunits used to stimulate S-specific CD4+ or CD8+ T cells.

| CD4 peptides | Subunit | Amino acid position |
|---|---|---|
| QDLFLPFFSNVTWFH | S1 | 52-66 |
| STEIYQAGSTPCNGV | S1 | 469-483 |
| VLSFELLHAPATVCG | S1 | 512-526 |
| ENSVAYSNNSIAIPT | S2 | 702-716 |
| ITSGWTFGAGAALQI | S2 | 882-896 |
| QMAYRFNGIGVTQNV | S2 | 901-915 |
| GKIQDSLSSTASALG | S2 | 932-946 |
| IRAAEIRASANLAAT | S2 | 1013-1027 |
| GYHLMSFPQSAPHGV | S2 | 1046-1060 |
| PAQEKNFTTAPAICH | S2 | 1069-1083 |
| **CD8 peptides** | **Subunit** | **Amino acid position** |
| FVFLVLLPL | S1 | 2-10 |
| VNLTTRTQL | S1 | 16-24 |
| LFLPFFSNV | S1 | 54-62 |
| SNVTWFHAI | S1 | 60-68 |
| VTWFHAIHV | S1 | 62-70 |
| RGWIFGTTL | S1 | 102-110 |
| FQFCNDPFL | S1 | 133-141 |
| YSSANNCTF | S1 | 160-168 |
| VSQPFLMDL | S1 | 171-179 |
| KIYSKHTPI | S1 | 202-210 |
| INITRFQTL | S1 | 233-241 |
| AAAYYVGYL | S1 | 262-270 |
| VRFPNITNL | S1 | 327-335 |
| FNATRFASV | S1 | 342-350 |
| GNYNYLYRL | S1 | 447-455 |
| VGYQPYRVV | S1 | 503-511 |
| VVVLSFELL | S1 | 510-518 |
| VNFNFNGLT | S1 | 539-547 |
| YQDVNCTEV | S1 | 612-620 |
| SIIAYTMSL | S2 | 691-699 |
| VAYSNNSIA | S2 | 705-713 |
| FGGFNFSQI | S2 | 797-805 |
| AALQIPFAM | S2 | 892-900 |
| VVNQNAQAL | S2 | 951-959 |
| VVFLHVTYV | S2 | 1060-1068 |
| ISGINASVV | S2 | 1169-1177 |
| IWLGFIAGL | S2 | 1216-1224 |
| IAIVMVTIM | S2 | 1225-1233 |

### S.Table 2

**Supplementary Table 2.** Primers used for construction and sequencing of SF-dER, S2-dER and their 2P mutation counterparts along with the primers used for mouse-adapted SARS-CoV-2 vRNA detection.

| Primer name | Sequence |
|---|---|
| **Construction primers** | |
| BsiWI-Signal | TAACGTACGGCCACCATGTTCGTCTTTCTGGTATTG |
| BssHII-SF | TAAGCGCGCCTATTATTCGGAATCATCCTCATCGA |
| BsmBI-signal | TAACGTCTCCGCACTGGGAACTCACCAGAGGAAG |
| BsmBI-S2 | TAACGTCTCAGTGCGTAGCAAGTCAGAGTATCATAG |
| BssHII-S2 | TAAGCGCGCTTATTCGGAATCATCCTCATCGAATTTAC |
| BsmBI-2P-fwd | AATCGTCTCACACCAGAGGCCGAAGTGCAGATTGATCGCCTG |
| BsmBI-2P-rev | ATACGTCTCAGGTGGGTCGAGCCGAGACAAGATGTCGTTC |
| **Sequencing primers** | |
| Signal-fwd1 | TCTGGTATTGCTTCCTCTGGTG |
| SF-fwd2 | TGCGCACTTGATCCATTGTC |
| S2-fwd3 | GTAAAGCACACTTCCCAAGAG |
| SF-fwd4 | GATCCTGGACATCACTCCATGC |
| SF-rev1 | TTCCACTTACATGGATAGCGTGG |
| S2-rev1 | TACTGTTATTACTATAAGCGACAG |
| S2-rev3 | GATCTCTAGCGGCGATATCTC |
| 3433 | GACCTTGGGAGGCAATCACT |
| oligo8a | GGAATCGCTGTCCTCAACAA |
| 9119 | AGATAGGGCTGCTAGTGAACCAAT |
| 9218 | TGGACCCTACGTTTTTCTTAATTCT |
| **qRT-PCR primers** | |
| nCoV_IP4-14059Fw | GGTAACTGGTATGATTTCG |
| nCoV_IP4-14146Rv | CTGGTCAAGGTTAATATAGG |
| nCoV_IP4-14084Probe(+) | TCATACAAACCACGCCAGG [5']Hex [3']BHQ-1 19 |
